# Essential roles of Hdac1 and 2 in lineage development and genome-wide DNA methylation during mouse preimplantation development

**DOI:** 10.1101/662767

**Authors:** Panpan Zhao, Huanan Wang, Han Wang, Yanna Dang, Lei Luo, Shuang Li, Yan Shi, Lefeng Wang, Shaohua Wang, Jesse Mager, Kun Zhang

## Abstract

Epigenetic modifications, including DNA methylation and histone modifications, are reprogrammed considerably following fertilization during mammalian early embryonic development. Incomplete epigenetic reprogramming is a major factor leading to poor developmental outcome in embryos generated by assisted reproductive technologies, such as somatic cell nuclear transfer. However, the role of histone modifications in preimplantation development is poorly understood. Here, we show that co-knockdown (cKD) of *Hdac1* and *2* (but not individually) resulted in developmental failure during the morula to blastocyst transition. This outcome was also confirmed with the use of small-molecule Hdac1/2-specific inhibitor FK228. We observed reduced cell proliferation and increased incidence of apoptosis in cKD embryos, which were likely caused by increased acetylation of Trp53. Importantly, both RNA-seq and immunostaining analysis revealed a failure of lineage specification to generate trophectoderm and pluripotent cells. Among many gene expression changes, a substantial decrease of *Cdx2* may be partly accounted for by the aberrant Hippo pathway occurring in cKD embryos. In addition, we observed an increase in global DNA methylation, consistent with increased DNA methyltransferases and Uhrf1. Interestingly, deficiency of Rbbp4 and 7 (both are core components of several Hdac1/2-containing epigenetic complexes) results in similar phenotypes as those of cKD embryos. Overall, Hdac1 and 2 play redundant functions required for lineage specification, cell viability and accurate global DNA methylation, each contributing to critical developmental programs safeguarding a successful preimplantation development.

**Significance:** Substantial changes to epigenetic modifications occur during preimplantation development and can be detrimental when reprogrammed incompletely. However, little is known about the role of histone modifications in early development. Co-knockdown of Hdac1 and 2, but not individually, resulted in developmental arrest during morula to blastocyst transition, which was accompanied by reduced cell number per embryo and increased incidence of apoptosis. Additionally, we observed a failure of first lineage specification to generate trophectoderm and pluripotent cells, which were associated with reduced expression of key lineage-specific genes and aberrant Hippo pathway. Moreover, an increase in global DNA methylation was found with upregulated Dnmts and Uhrf1. Thus, Hdac1 and 2 play overlapping roles in lineage development, apoptosis, and global methylation during preimplantation development.

## Introduction

A distinguishing feature of preimplantation development is a remarkable reprograming of the epigenome, including DNA modifications and post-translational histone modifications (1, 2). Aberrant epigenetic reprograming has been associated with defects in various biological processes, including DNA replication and embryonic genome activation (EGA), which eventually leads to early embryonic death (3). Moreover, incomplete epigenetic reprogramming is a major contributing factor to the poor developmental outcome associated with the use of assisted reproductive technologies, including in vitro embryo production (IVP) (4) and somatic cell nuclear transfer (SCNT) (5–10). Indeed, modulation of certain epigenetic modifications has been proved a viable tool to enhance SCNT rate and obtain live cloned monkeys (11). However, little is known about the epigenetic regulation of critical developmental events (e.g. lineage development) and interactions between epigenetic modifications in preimplantation embryos.

Histone deacetylase (Hdac) 1 and 2 are highly homologous enzymes present together in multiprotein complexes, the most extensively characterized being NuRD (12), Sin3 (13), and CoREST (14), which are conserved ranging from yeast to human (15, 16). Histone acetylation is well known for its role in transcriptional activation through opening of chromatin and nucleosome compaction (17). Accordingly, Hdac1/2-containing complexes are traditionally thought to act as transcriptional corepressors of target genes. However, Hdac1/2-containing complexes have also been shown to be tethered to actively transcribed genes, suggesting a critical role in transcriptional activation in certain situations (15, 16, 18, 19).

Because of high homology and physical colocalization in large multiprotein complexes, it is reasonable that Hdac1 and Hdac2 are functionally redundant in multiple biological systems (20–23). However, specific roles of Hdac1 and 2 have also been documented. For instances, *Hdac2* is specifically involved in the regulation of memory formation and synaptic plasticity (24). In contrast, knockout of *Hdac1* results in early lethality at peri-implantation stage (25). Furthermore, knockdown of both maternal and zygotic *Hdac1* or *Hdac2* by siRNA injection in preimplantation embryos results in no difference on blastocyst formation (26). These relatively mild phenotypes in preimplantation embryos may be caused by the functional redundancy of *Hdac1* and *2*. Therefore, the precise role of Hdac1 and 2 and the underlying molecular mechanisms during preimplantation embryogenesis remain unresolved.

In this study, we show that double knockdown of *Hdac1* and *2*, but not individually, resulted in lethality during the morula to blastocyst transition. The developmental failure is accompanied by a substantial perturbation of the transcriptomes and lineage development in conjunction with increased incidence of apoptosis, enhanced histone acetylation and genome-wide DNA methylation. We propose that Hdac1 and 2 play compensatory and essential roles during preimplantation development, at least partly through modulation of lineage specification, apoptosis and global DNA methylation.

## Results

### Double knockdown of *Hdac1* and *2* results in developmental arrest during morula to blastocyst transition

Previous RNA-seq (GSE44183, Fig. S1*A*) and quantitative PCR analysis revealed extensive expression of *Hdac1* and *2* through preimplantation development (27, 28). As anticipated, we confirmed Hdac1 and 2 concentrated and co-localized in nucleoplasm of blastomeres from 2-cell to blastocyst stage (Fig. S1*B*). Moreover, both proteins appear evenly distributed in trophectoderm cells (TE) and inner cells mass (ICM) in the mouse blastocyst (Fig. S1*B*). Overall, these results imply Hdac1 and 2 may play an overlapping role during preimplantation development.

Both *Hdac1* and *2* are maternally derived prior to the occurrence of the major wave of EGA at 2-cell in mice (22, 28). Conditional double knockout of *Hdac1* and *2* in oocytes results in oogenesis failure (22), prohibiting us from establishing knockout models of both maternal and zygotic *Hdac1* and *2*. We therefore decided to employ two complementary approaches to investigate the *in vivo* roles for *Hdac1* and *2* in preimplantation development: RNAi and Hdac1/2-specific small-molecule inhibitor, FK228 (29) (Fig. 1*A*).

**Fig. 1.**
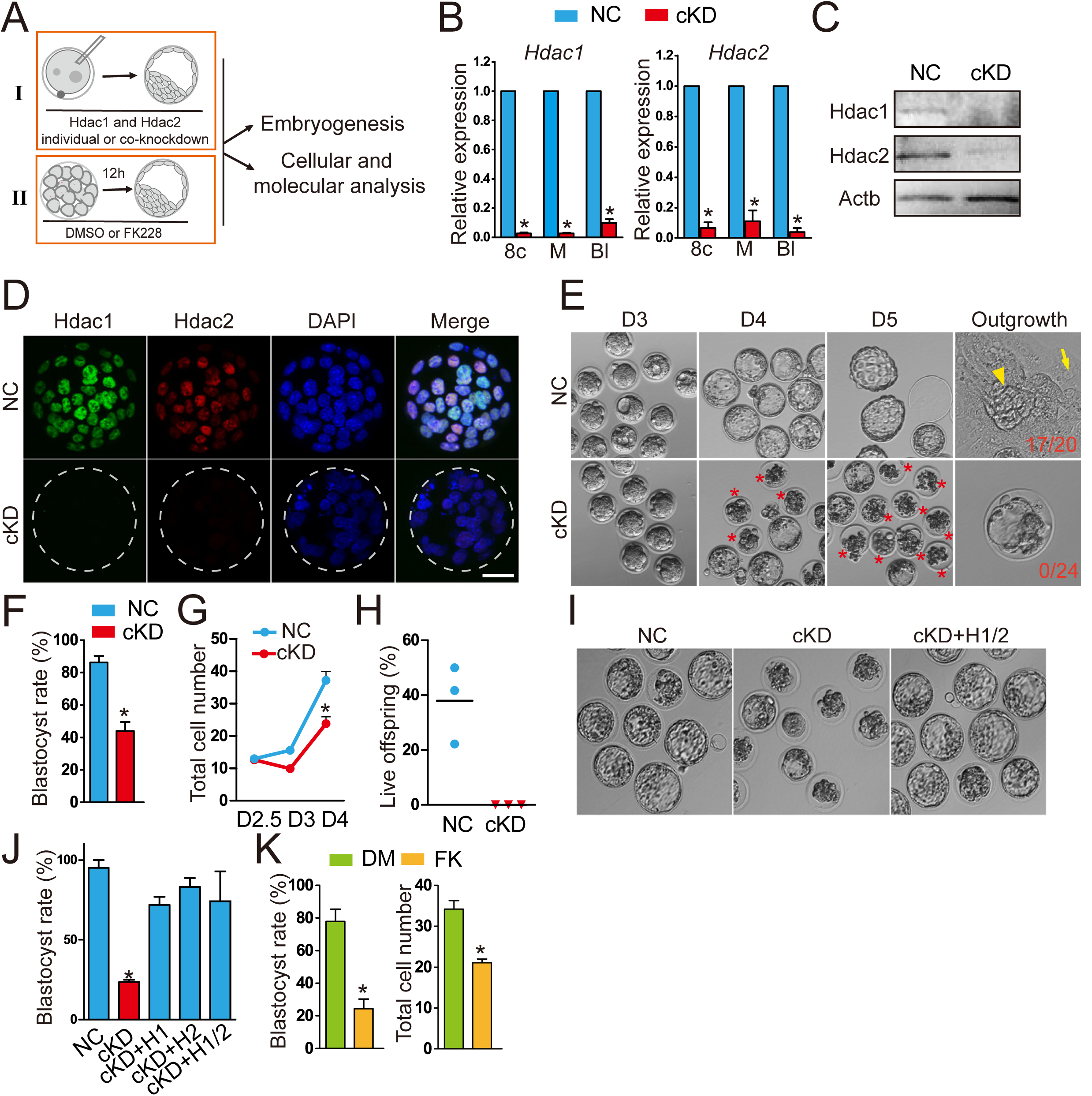
Double knockdown of Hdac1 and 2 results in embryonic lethality during the morula to blastocyst transition. (A) Schematic overview of two approaches (I: RNAi; II: small-molecule inhibitor, FK228) used to investigate *in vivo* roles of Hdac1 and 2 during preimplantation development. (B) qPCR analysis of knockdown efficiency of siRNA cocktails targeting Hdac1 and 2 from 8-cell to blastocyst stage. Mouse zygotes derived *in vivo* were microinjected with Hdac1/2 siRNA cocktails (20 µM, 10 pl, cKD) or negative control siRNAs (NC). Embryos were collected at 8-cell (8c), morula (M) and blastocyst (Bl) stage (n=3 pools of 5-10 embryos each per treatment). Data were stated as mean ± SEM normalized to endogenous control (*H2afz*; *P<0.05). (C) Immunoblot analysis of Hdac1 and 2 in NC and cKD morulae (30 embryos per group, 2 replicates were performed with similar results). β-actin (Actb) was used as a loading control. (D) Immunocytochemical detection of dramatic reduction of Hdac1 and 2 protein in cKD blastocysts. Three replicates were conducted and at least 10 embryos analyzed in each group (Scale bar: 25 µm). (E) Representative photos of NC and cKD embryos from Day 3 after mating (D3) to D5. Arrow head: ICM outgrowth; Arrow: trophoblast giant cell. Asterisk: Degenerated embryos. (F) Blastocyst rate in NC and cKD groups at D4 (n=5; 16-33 embryos per group per replicate). Data are shown as mean ± SEM (*P<0.05). (G) Cell counting analysis of NC and cKD embryos from D2.5 to D4 (n=3). (H) Percent live offspring out of embryos transferred (n=3; 15-20 embryos were transferred per group). (I and J) Rescue of cKD embryos by microinjection of exogenous *Hdac1* and/or *Hdac2* mRNA (n=3; 15-20 embryos per group; *P<0.05). (K) Blastocyst rate and total cell number per embryo in embryos treated with Hdac1/2 specific inhibitor, FK228 (n=3; 15-20 embryos per group). Data are expressed as mean ± SEM. Different superscripts indicate significant differences (P < 0.05).

The effectiveness of the siRNAs and the inhibitor was verified. Analysis of qPCR revealed *Hdac1* mRNA level was depleted by approximately 90% from 8-cell to blastocyst stage (n=3, P<0.05) after microinjection of *Hdac1* siRNA cocktail (H1 KD) relative to control embryos injected with nonspecific siRNA (NC, Fig. S2*A*). In accordance, Hdac1 protein abundance was also depleted (n=3, Fig. S2*B*). Hdac2 protein abundance was not affected by H1 KD, suggesting a robust specificity of the siRNA (Fig. S2*B*). Similarly, the *Hdac2* siRNA cocktail (H2 KD) produced a 90% knockdown at the mRNA level (n=3, P<0.05; Fig. S2*A* and 2*B*). Co-microinjection of siRNAs targeting *Hdac1* and *2* (cKD) resulted in dramatic decreases in both endogenous *Hdac1* (above 90% reduction) and *Hdac2* (above 89% reduction) between the 8-cell to blastocyst stage (n=3, P<0.05; Fig. 1*B*). Immunoblotting (n=2) and IF (n=3) analysis confirmed a successful reduction of the amount of Hdac1 and 2 protein in morula and blastocysts (Fig. 1*C* and 1*D*). Because of Hdac1 and 2’s critical roles in histone de-acetylation, IF was performed to determine if histone acetylation was affected. The amount of histone H3 lysine 14 acetylation (H3K14ac) and H4K5ac was increased by 72.9% and 64.4%, respectively, in cKD embryos relative to controls (n=3, P<0.05; Fig. S3*A* and 3*B*). Similarly, treatment of mouse morula with FK228 increased both H3K14ac (by 116%) and H4K5ac (by 67.0%) relative to the vehicle control (DMSO; n=3, P<0.05; Fig. S3*D* and 3*E*). Taken together, these results suggest the siRNAs and inhibitor are highly effective in the context of preimplantation development.

We next monitored the developmental potential of embryos of NC, H1 KD, H2 KD, and cKD. No morphological difference was observed in H1 KD and H2 KD groups compared with NC throughout preimplantation development (Fig. S4*A*), consistent with a previous study (26). Both the blastocyst rate (above 80%, Fig. S4*B*) and total cell number per embryo (Fig. S4*C*) were normal in the H1 KD and H2 KD groups. By contrast, the cKD embryos appeared normal up to the morula stage (Fig. 1*E*) but more than half of cKD embryos fail to develop into blastocysts (n=5, P<0.05; Fig. 1*F*). Cell counting analysis revealed that total cell number per embryo declined from D3 (D3: 15.6±1.1 vs 9.9±0.9; D4: 37.2±2.8 vs 23.8±2.1; P<0.05; Fig. 1*G*). To evaluate if the development of cKD embryos was delayed, the embryos were continually cultured *in vitro* until D5. At D5, the majority of cKD embryos collapsed whereas NC blastocysts completed hatching, ruling out the possibility of developmental delay (Fig. 1*E*). To test the developmental competency of the blastocysts that do develop in cKD groups, blastocysts were cultured individually to examine if outgrowths could be formed. Consistent with our previous report (30), the potential to form embryo outgrowth is compromised in H1 KD group, but not H2 KD group (Fig. S4*D*). In contrast, none of cKD blastocysts are capable to form outgrowths (0/24 vs 17/20, n=3), even after zona pellucida removal (Fig. 1*E*). To test if the *in vivo* environment could alleviate the phenotype, 2-cell embryos of NC and cKD groups were transferred into surrogates. Embryo transfer analysis revealed that cKD embryos failed to generate live offspring whereas 22-50% embryos transferred in NC group developed to term (n=3, Fig. 1*H*). In sum, these results suggest Hdac1 and 2 play a compensatory role in supporting preimplantation development.

Three experiments were performed to verify the initial siRNA findings. First, an alternative cocktail of siRNAs targeting 5’ and 3’ untranslated regions (5’ and 3’ UTR) of *Hdac1* and *2* was used. Developmental arrest during morula to blastocyst transition was also observed with reduced blastocyst rate (23.6% vs 95.0% in NC; n=3, P<0.05; Fig. 1*I* and 1*J*). Second, the development of cKD embryos could be rescued (blastocyst rate>70%) by co-injection of exogenous *Hdac1* and/or *Hdac2* mRNA transcribed *in vitro* that were not targeted by the siRNAs (n=3, P<0.05; Fig. 1*I*, *1J*, S4*E*, and S4*F*). Last, treatment of mouse morula with FK228 also resulted in reduced blastocyst rate (24.3% vs 77.9% in control group, n=3, P<0.05) and total cell number per embryo (34 vs 21, n=3; Fig. 1*K*). Overall, a combination of loss of function approaches (RNAi plus small-molecule inhibitor) and rescue experiments confirm the specificity of our approach and the essential role of Hdac1 and 2 in preimplantation development.

### Effect of cKD on transcriptomic profile of preimplantation embryo

To delineate the molecular basis of the developmental arrest of cKD embryos, we carried out RNA-seq in NC and cKD morulae, obtained prior to the emergence of morphological phenotypes (to avoid bias, Fig. 2*A*). Hierarchical clustering revealed a separation between NC and cKD morulae (n=3; Fig. S5*A*). We found that 991 genes were differentially expressed (Fold changes (FC) >2 or <0.5, P adjusted<0.05; Table S1), 72% of which were upregulated, consistent with the notable role of Hdac1/2 as transcriptional repressors. Expression of select genes (Down: *Myc*, *Dab2*, *Amot*, *Fgfr2*, and *Otx2*; Up: *Arid3a* and *Sfmbt2*; No change: *Tet1* and *Ctnnb1*) was confirmed by qPCR analysis (Fig. 2*B*).

**Figure 2.**
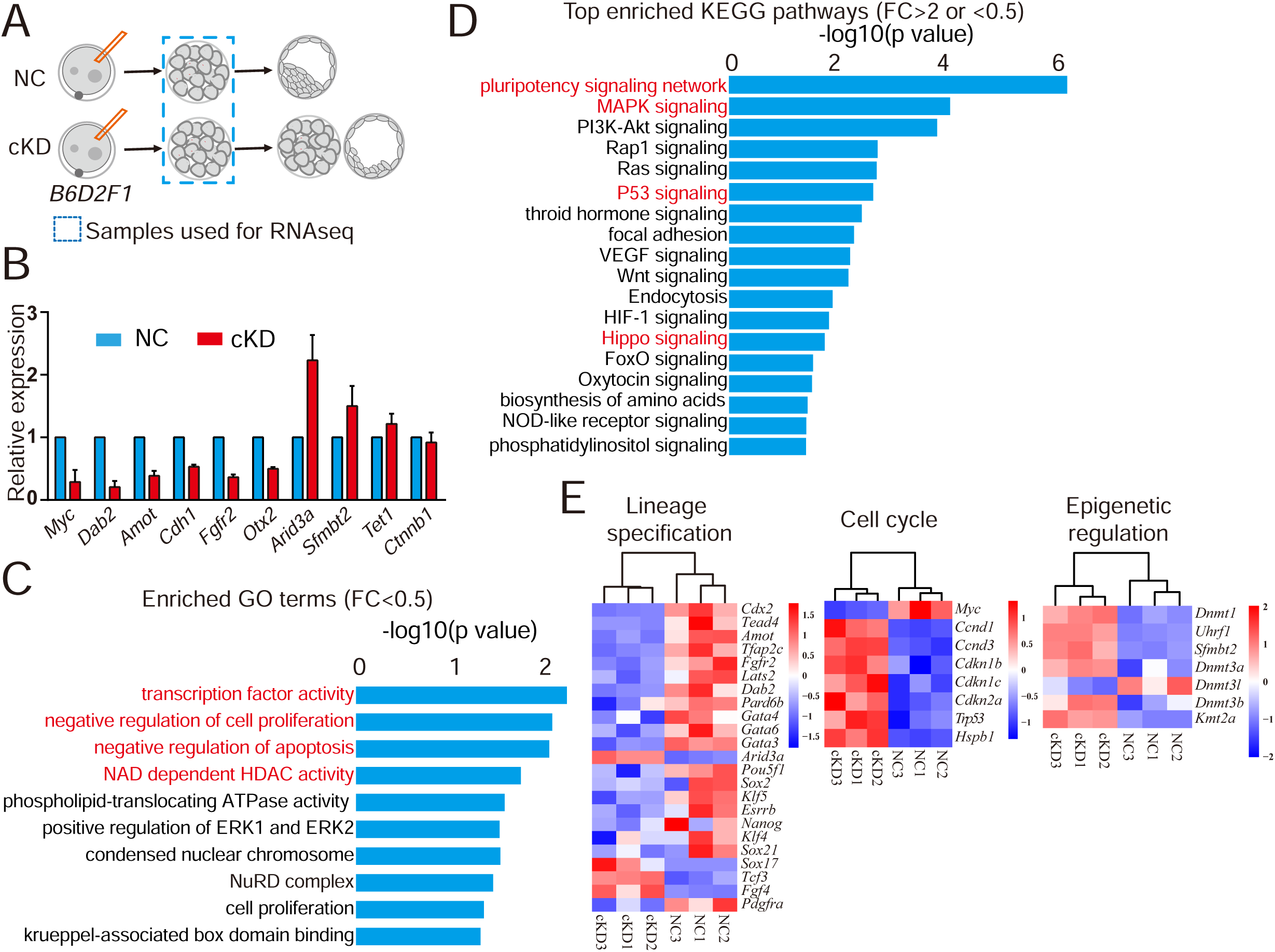
RNA-seq analysis of embryos deficient of Hdac1 and 2. (A) Schematic overview of the samples collected for RNA-seq analysis (n=3; 60 embryos/group/replicate). (B) Validation of RNA-seq results on expression levels of selected genes (Downregulated:*Myc*, *Dab2*, *Amot*, *Cdh1*, *Fgfr2*, *Otx2*; Upregulated: *Arid3a*, *Sfmbt2*; No change: *Tet1*, *Ctnnb1*). Three biological replicates were performed with 5-10 morula collected for each group (*P<0.05). (C) GO analysis of downregulated genes in cKD morulae. The data indicate enriched GO terms related to epigenetic regulation, cell proliferation and apoptosis. (D) KEGG analysis of differentially expressed genes (DEGs) between NC and cKD morulae. The data indicate cKD leads to abnormal signaling pathway of pluripotency network, P53 and Hippo. (E) Overrepresentation of genes related to lineage specification, cell cycle and epigenetic regulation among DEGs.

Gene ontology (GO) analysis revealed that the top GO terms (biological processes) enriched in differentially expressed genes (DEGs; FC>2 or <0.5) include processes involved in DNA transcription, cell differentiation, cell proliferation and apoptosis (Fig. S5*B*). Specifically, GO analysis of downregulated genes (FC<0.5) showed that enriched GO terms include transcription factor activity, cell proliferation, apoptosis, NAD dependent Hdac acitivity and NuRD complex (Fig. 2*C*). Moreover, KEGG analyses revealed top hits in signaling pathways regulating pluripotency, MAPK, P53 and Hippo signaling (Fig. 2*E*).

Among the downregulated genes, we observed an over-representation of TE specific genes (*Cdx2*, *Dab2, Fgfr2*), genes associated with Hippo signaling (*Tead4, Amot, Lats2*) and genes related with pluripotency networks (*Nanog*, *Klf5*, *Sox2*, *Pou5f1, Myc*) (Fig. 2*C*). Among the upregulated genes, we observed an enrichment of genes related to cell cycle progression and apoptosis (*Trp53*, *Ccnd1, Ccnd3, Cdkn1b, Cdkn1c, Cdkn2a*) and genes related to chromatin modification, including *Dnmt1* and *Uhrf1* (Fig. 2*C*). Taken together, the transcriptome profiling implies that embryos lacking Hdac1 and 2 do not properly initiate early lineage differentiation, cell proliferation, and genome-wide methylation in preimplantation embryos.

### Hdac1 and 2 reduction leads to increased apoptosis, increased Trp53 acetylation and defective proliferation in preimplantation embryos

Because total cell number per embryo was drastically reduced in cKD (Fig. 1*G*) and RNA-seq analysis in cKD embryos identified genes related with apoptosis among the DEGs (Fig. 2*D*), we performed assays to test if apoptosis was abnormal in cKD embryos. The incidence of apoptosis was markedly increased in cKD blastocysts (90%, n=10) relative to controls (10%, n=10) (Fig. 3*A*).

**Figure 3.**
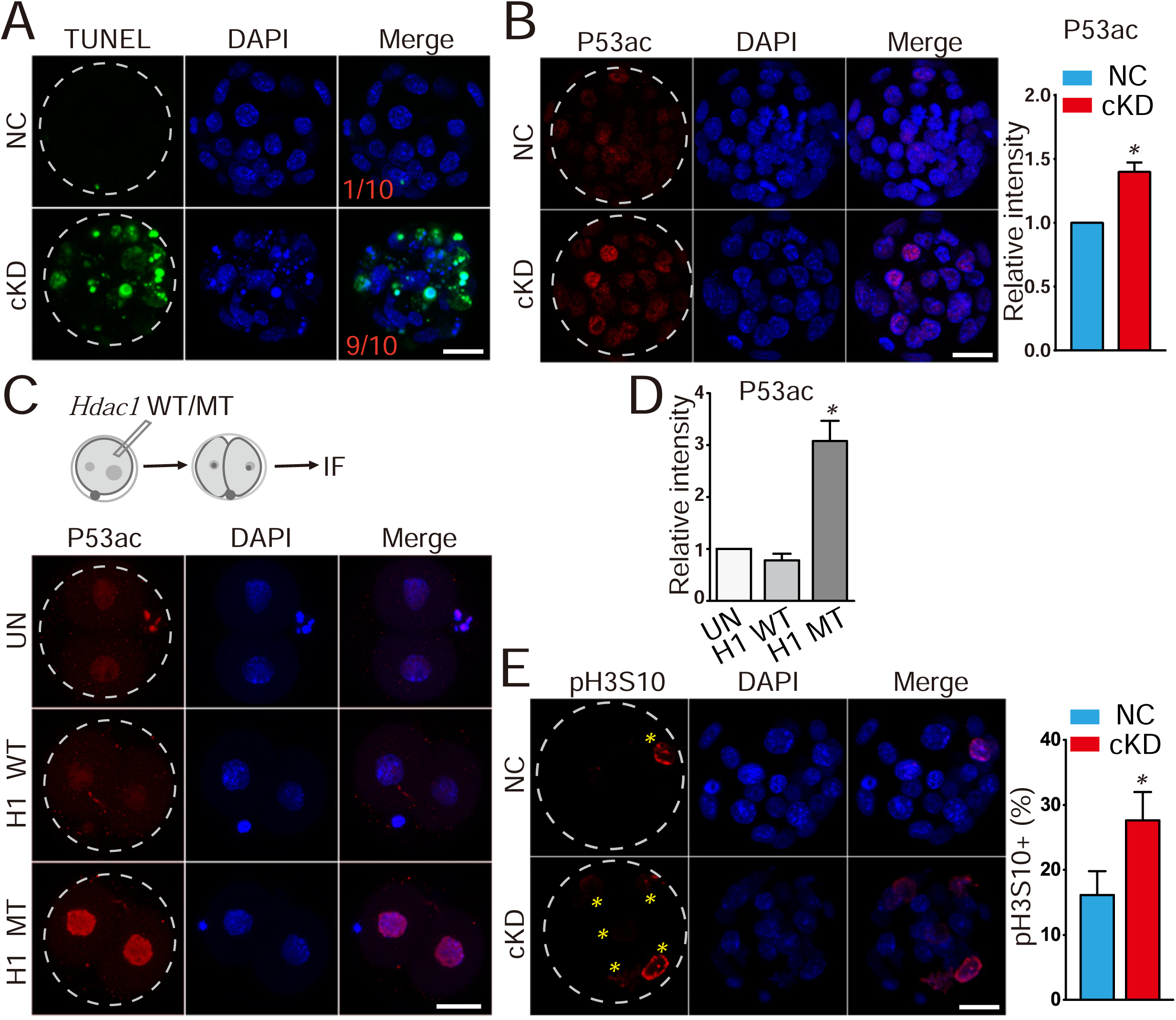
Hdac1 and 2 deficiency leads to increased incidence of apoptosis, increased Trp53 acetylation and cell proliferation arrest. (A) TUNEL analysis of NC (n=10) and cKD blastocysts (n=10). The data revealed a dramatic increase of incidence of apoptosis in cKD blastocysts. Three biological replicates were conducted. (B) Immunocytochemical analysis of Trp53 acetylated on K379 (P53ac) in blastocysts. The intensity of P53ac was improved significantly (n=3; 5-10 embryos per group per replicate, Scale bar: 25 μm). Nuclear was counterstained with DAPI. (C) *Hdac1* was mutated at the deacetylase site and mRNA was *in vitro* produced. Wildtype *Hdac1* (H1 WT) and mutant *Hdac1* (H1 MT) was introduced into zygote and 2-cell embryos were collected for immunocytochemical analysis (n=3; 5-10 embryos per group per replicate, Scale bar: 25 μm). (D) The intensity of P53ac was not changed in H1 WT embryos but increased in H1 MT embryos (*P<0.05). (E) Immunocytochemical examination of histone H3 serine 10 phosphorylation (pH3S10), a marker for late G2 and mitosis, in blastocysts (n=3; 5-10 embryos per group per replicate; Asterisk: pH3S10 positive blastomere; Scale bar: 25 μm).

Trp53 is a critical molecule regulating apoptosis and was also upregulated in cKD morulae as determined by RNA-seq. In addition, Trp53 activity has been shown to be repressed in an Hdac1-dependent manner through de-acetylation (31). Our results showed that the amount of Trp53 acetylation at lysine 379 (p53ac) was greater in cKD (Fig. 3*B*) or FK228-treated embryos relative to controls (Fig. S6*A*). To ascertain if Hdac1’s deacetylase activity is directly responsible for Trp53 acetylation in the context of preimplantation development, we performed mutagenesis at the deacetylase site of *Hdac1* and injected wild-type *Hdac1* (H1 WT) and mutant *Hdac1* (H1 MUT) mRNA into zygotes (Fig. 3*C*). No difference was observed in H1 WT-injected embryos relative to uninjected controls (UN; Fig. 3*C* and 3*D*). In contrast, there is a dramatic increase of p53ac in H1 MUT-injected embryos (Fig. 3*D*). Overall, these results indicate Hdac1’s enzymatic activity is directly responsible for deacetylation of the non-histone protein, Trp53, during preimplantation stages.

To ascertain if cell proliferation was affected by cKD, we performed IF against histone H3 serine 10 phosphorylation (pH3S10), a marker for late G2/M phase. Only 16.2% of blastomeres in control morulae were subject to mitosis whereas the incidence of pH3S10 positive blastomeres was increased significantly in cKD embryos (27.7%; Fig. 3*E*), suggesting a cell cycle block at G2/M phase.

Interphase bridges have recently been identified as a critical subcellular structure for mouse preimplantation embryos (32). As anticipated, interphase bridges were detected at cell-cell junctions in control embryos (Fig. S6*B* arrows). Number of interphase bridges in H1 KD or H2 KD embryos is comparable or increased relative to controls but was reduced in cKD embryos (Fig. S6*B*), suggesting an aberrant cellular communication in the absence of Hdac1/2.

### Double knockdown of Hdac1 and 2 results in failed lineage specification of trophectoderm and inner cell mass

Transcriptome profiling revealed substantial enrichment of TE-specific and pluripotency network genes among DEGs (Fig. 2*C*). The earliest lineage specification takes place during the morula to blastocyst transition and generate TE (precursors of the majority of placental cells) and ICM (precursors of the embryo proper), we thus decided to examine the cell differentiation program in cKD embryos. We quantified the expression of Cdx2, a critical molecular marker of TE. Abundance of *Cdx2* mRNA was unchanged through blastocyst stage in H2 KD embryos and downregulated slightly in H1 KD morulae and blastocysts (Fig. S7*A*). IF results displayed a normal distribution of Cdx2 signal in both H1 and H2 KD embryos (Fig. S7*B*). In contrast, Cdx2 mRNA and protein were diminished in cKD embryos during the morula to blastocyst transition (Fig. 4*A*-*C*), which was confirmed in FK228-treated embryos (Fig. 4*D* and 4*E*). The expression of *Cdx2* in cKD embryos could be successfully rescued by injection of either *Hdac1* or *2* mRNA (Fig. 4*F* and 4*G*). To further determine if reduced expression of Cdx2 was cell-autonomous, we injected siRNAs into one of two blastomeres at 2-cell stage and *H2B-RFP* was used as a lineage-tracing marker (Fig. 4*H*). Surprisingly, we found Cdx2 disappeared not only in blastomeres derived from siRNA-injected but un-injected cells, suggesting Hdac1/2 is involved in regulation of signaling molecules upstream of *Cdx2* expression (Fig. 4*H*).

**Figure 4.**
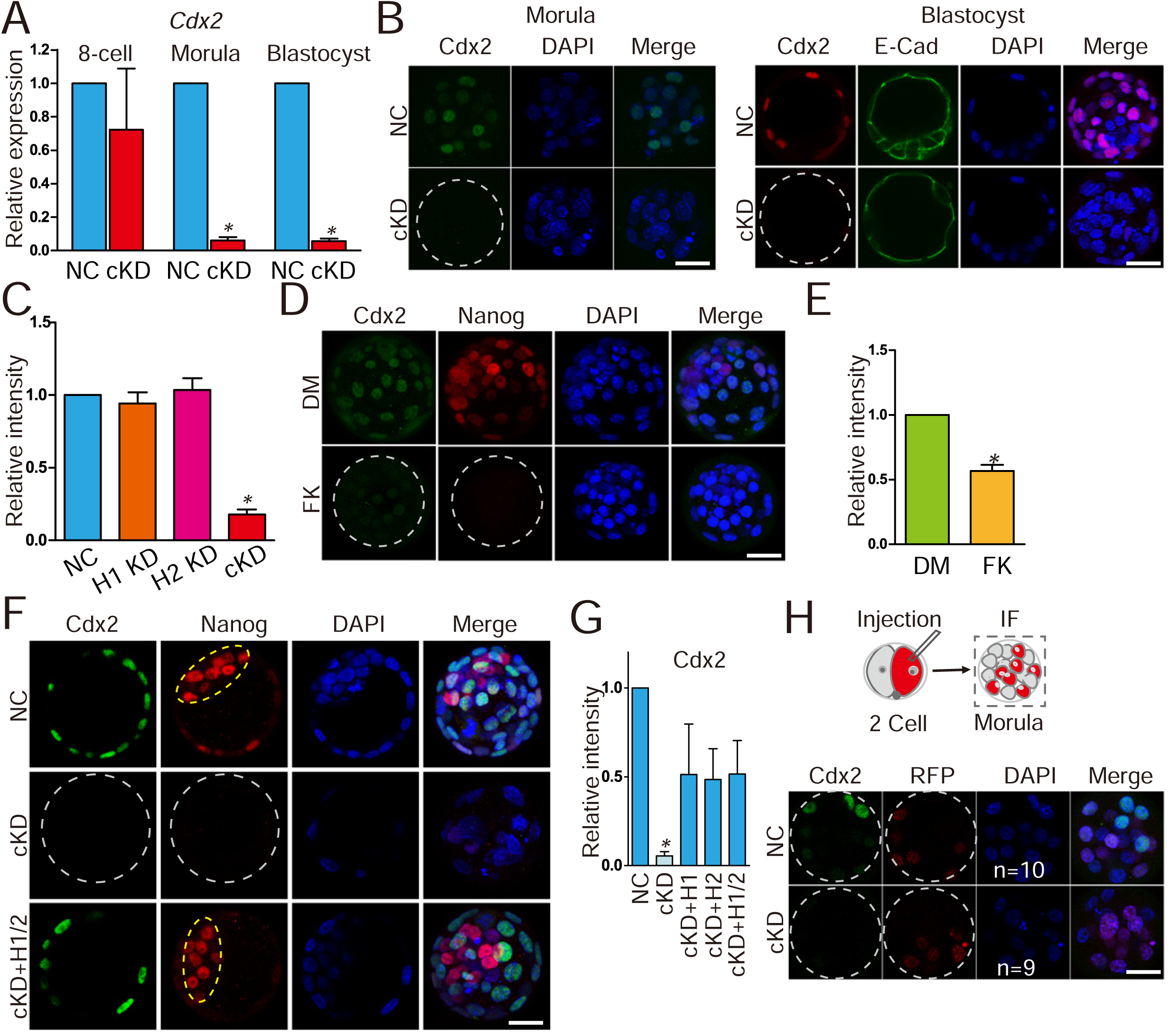
Cdx2 was inactivated in embryos deficient of Hdac1 and 2. (A) qPCR analysis of *Cdx2* in 8-cell embryos, morula and blastocysts (n=3 pools of 5-10 embryos each per group). (B-E) Immunocytochemical analysis of Cdx2 in morula and blastocysts after RNAi (B and C) or FK228 treatment (D and E). Three biological replicates with 5-10 embryos analyzed per group each time. The intensity of Cdx2 was diminished in cKD and FK228 treated, but not H1 or H2 KD embryos (C and E). E-Cad: E-Cadherin; DM: DMSO; FK: FK228. (F and G) Rescue of Cdx2 in cKD embryos after injection of exogenous *Hdac1* or *2*. The experiment was conducted three times and 5-10 embryos analyzed per group per time. Yellow dashed oval: inner cell mass. (H) Nonspecific siRNAs or siRNAs cocktail targeting *Hdac1* and *2* were microinjected into one blastomere at 2-cell stage. *H2B-RFP* mRNA was co-injected as a tracking marker. Blastocysts were collected for immunocytochemical analysis (n = 3; 5-10 embryos per group per replicate). The intensity of Cdx2 was diminished not only in cells derived from the siRNA-injected blastomere but those from noninjected blastomere in cKD groups.

We next examined if the molecular signature of ICM was disrupted in embryos lacking *Hdac1* and *2*. Expression of Oct4, Nanog and Sox2 at both mRNA and protein level was unchanged in H1 or H2 KD embryos (Fig. S8*A* and 8*B*). However, mRNA level of *Oct4*, *Nanog* and *Sox2* was reduced in cKD groups during the morula to blastocyst transition (Fig. 5*A*). Similarly, FK228 treatment led to a decrease in mRNA abundance of *Oct4* and *Nanog* (Fig. S8*C*). IF results indicated no significant change of Oct4 signal, however, Nanog and Sox2 levels were dramatically decreased in cKD blastocysts (Fig. 5*B*-*C* and Fig. S8*E*), which was also confirmed using FK228 (Fig. S8*D*). Collectively, these data demonstrate a failure of the first cell fate decision that normally gives rise to TE and ICM cells.

**Figure 5.**
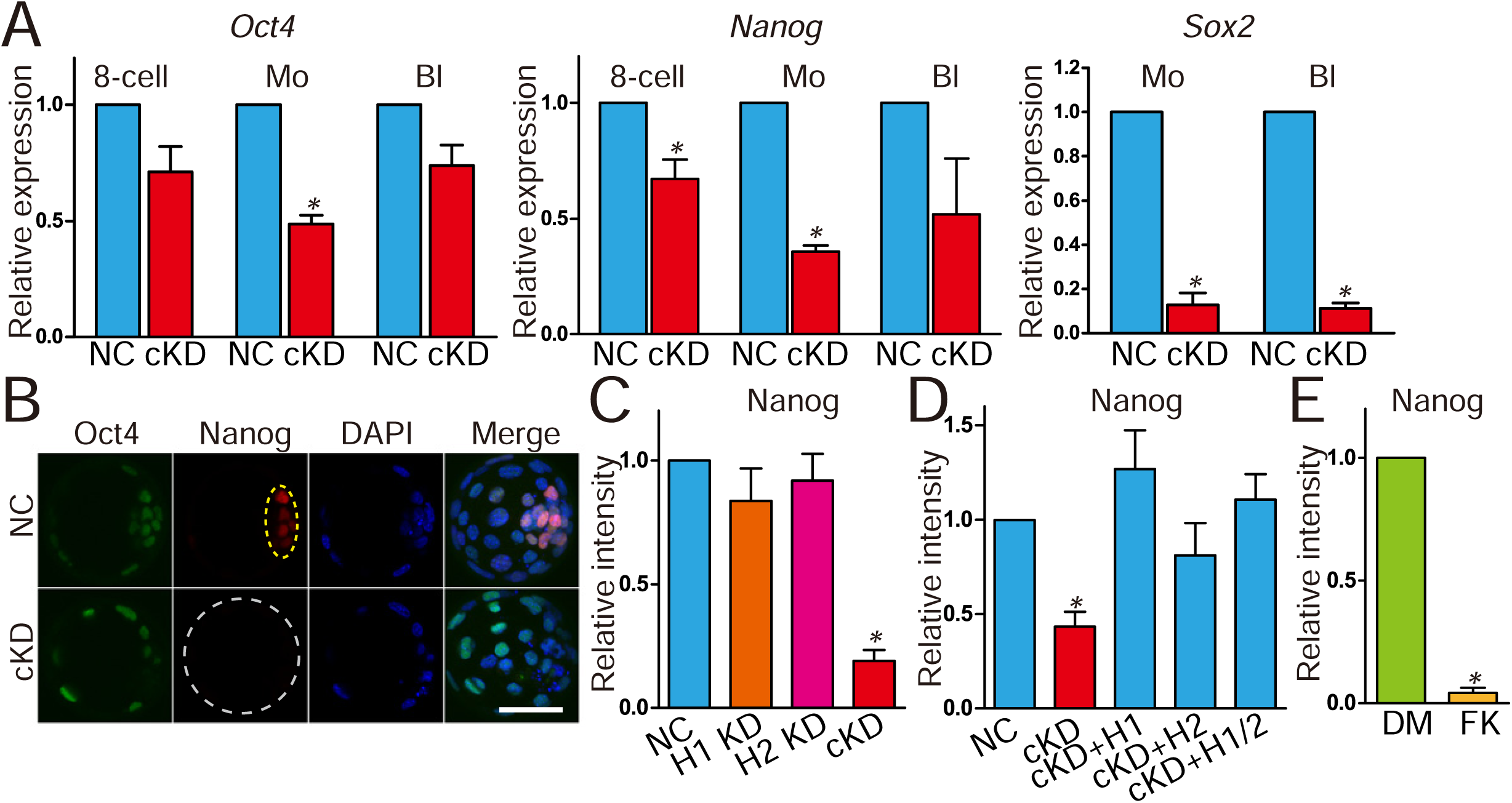
Key pluripotency genes Oct4, Nanog and Sox2 were downregulated in embryos deficient of Hdac1 and 2. (A) qPCR analysis of *Oct4, Nanog and Sox2* in NC and cKD embryos (n=3 pools of 5-10 embryos each per group). (B and C) Immunocytochemical analysis of Oct4 and Nanog in blastocysts after RNAi. The intensity of Nanog, but not Oct4 was diminished in cKD embryos (panel C; n=3; 5-10 embryos were analyzed per group each time, *P<0.05). Yellow dashed oval: inner cell mass. (D) Rescue of Nanog in cKD embryos after injection of exogenous *Hdac1* and/or *2*. The experiment was conducted three times and 5-10 embryos analyzed per group per time. (E) Analysis of the intensity of Nanog in embryos treated with either DMSO (DM, vehicle control) or FK228 (FK).

The second lineage specification occurs in the late blastocysts when the ICM differentiates into epiblast (Epi) and primitive endoderm (PrE). We examined Gata6, a marker of PrE, and Nanog, a marker of Epi, to determine if the second lineage specification failed as well. Results showed Gata6 and Nanog are mutually exclusively distributed in ICM in control blastocysts, however, no Gata6 and Nanog positive cells were visible in cKD embryos (Fig. S8*F*), confirming a failure of the earliest two lineage specification programs in mouse preimplantation embryos.

### Abnormal Hippo pathway in embryos deficient of both Hdac1 and 2

Hippo pathway components were enriched in GO analysis of DEGs between cKD and control morulae (Fig. 2*E*). Hippo pathway plays a critical role in defining TE specification program during mouse preimplantation development (33). Starting from the morula stage, Tead4 and Yap1 act as upstream regulators of Cdx2 and localize in the nucleus of TE cells (33). IF results showed that no visible difference was detected in Tead4 and Yap1 in H1 or H2 KD embryos (Fig S9*A*-*C*). However, *Tead4* mRNA was reduced in cKD from 8-cell to blastocyst stage while *Yap1* mRNA was slightly reduced (Fig 6*A*). IF analysis revealed that the number of Tead4 positive blastomeres was reduced by 50% in cKD group (Fig 6*B* and 6*C*). Immunoblotting analysis further confirmed that the protein abundance of Tead4 was diminished in cKD morulae (Fig 6*D*). Additionally, the percent Yap positive cells declined by 75% in cKD groups relative to controls (Fig 6*E* and 6*F*). Lats1 and Lats2 are upstream molecules that modulate the activity of Yap (34). Our qPCR results documented that mRNA of both genes was reduced significantly in cKD morulae (Fig S9*A*), suggesting their abnormal expression could account for defective Hippo signaling that we observed in the absence of Hdac1/2 activity.

**Figure 6.**
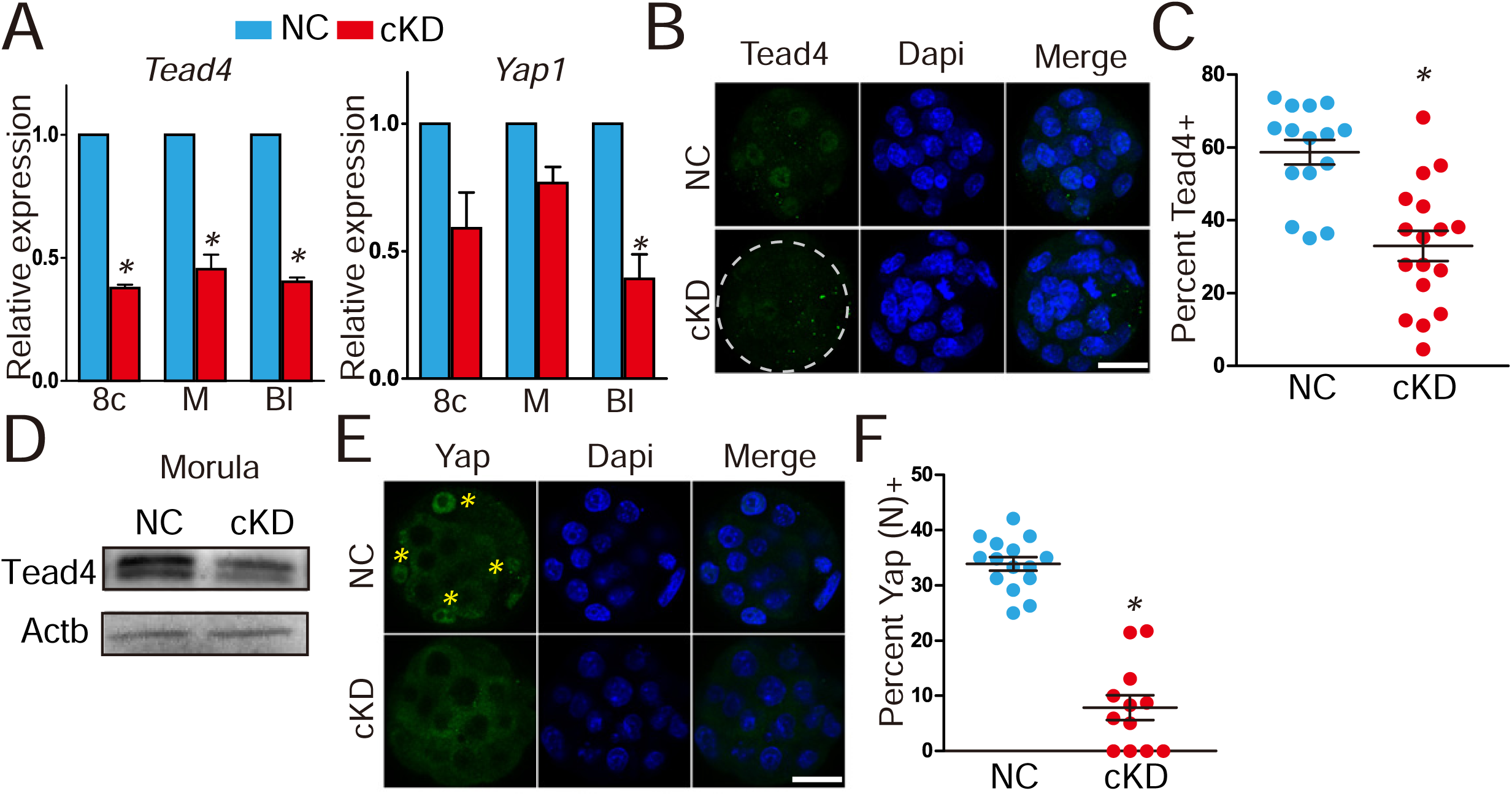
Hda1 and 2 co-knockdown results in aberrant Hippo signaling pathway. (A) qPCR analysis of *Tead4* and *Yap1* in 8-cell embryos (8c), morula (M) and blastocysts (Bl) (n=3 pools of 5-10 embryos each per group). (B-D) Immunocytochemical (n=3; 6-10 embryos were analyzed per group each time,, *P<0.05) and immunoblot analysis (n=2 pools of 30 embryos each per group, similar effects were obtained) of Tead4 in morula after RNAi. Both percent Tead4 positive cells (panel C) and the intensity of Tead4 (panel D) was reduced in cKD. (E and F) Immunocytochemical analysis of Yap in morula (n=3; 5-10 embryos were analyzed per group each time). Asterisk: nuclear Yap.

### Genome-wide DNA methylation was enhanced in blastocysts deficient of both Hdac1 and **2**

A wave of genome-wide DNA demethylation occurs after fertilization through preimplantation development, the molecular mechanism of which remains unclear (1). Changes in the expression of *Dnmt1* and *Uhrf1* were notable in our RNA-seq analysis given their central role in DNA methylation (Fig 2*E*). Thus, we sought to determine the global DNA methylation by examining 5-cytosine methylation (5mc) and 5-cytosine hydroxymethylation (5hmc), a newly defined DNA modification. Amounts of both DNA modifications are increased in cKD, but not in individual KD groups at blastocyst stage (n=3, P<0.05; Fig 7*A*), which was also seen in FK228-treated embryos (n=3, Fig S10*C*). However, little effect was observed on histone H3 lysine 4 trimethylation (H3K4me3), a marker for transcriptional activation, and histone H3 lysine 9 dimethylation (H3K9me2), a marker for transcriptional repression (Fig S10*A* and S10*B*).

**Figure 7.**
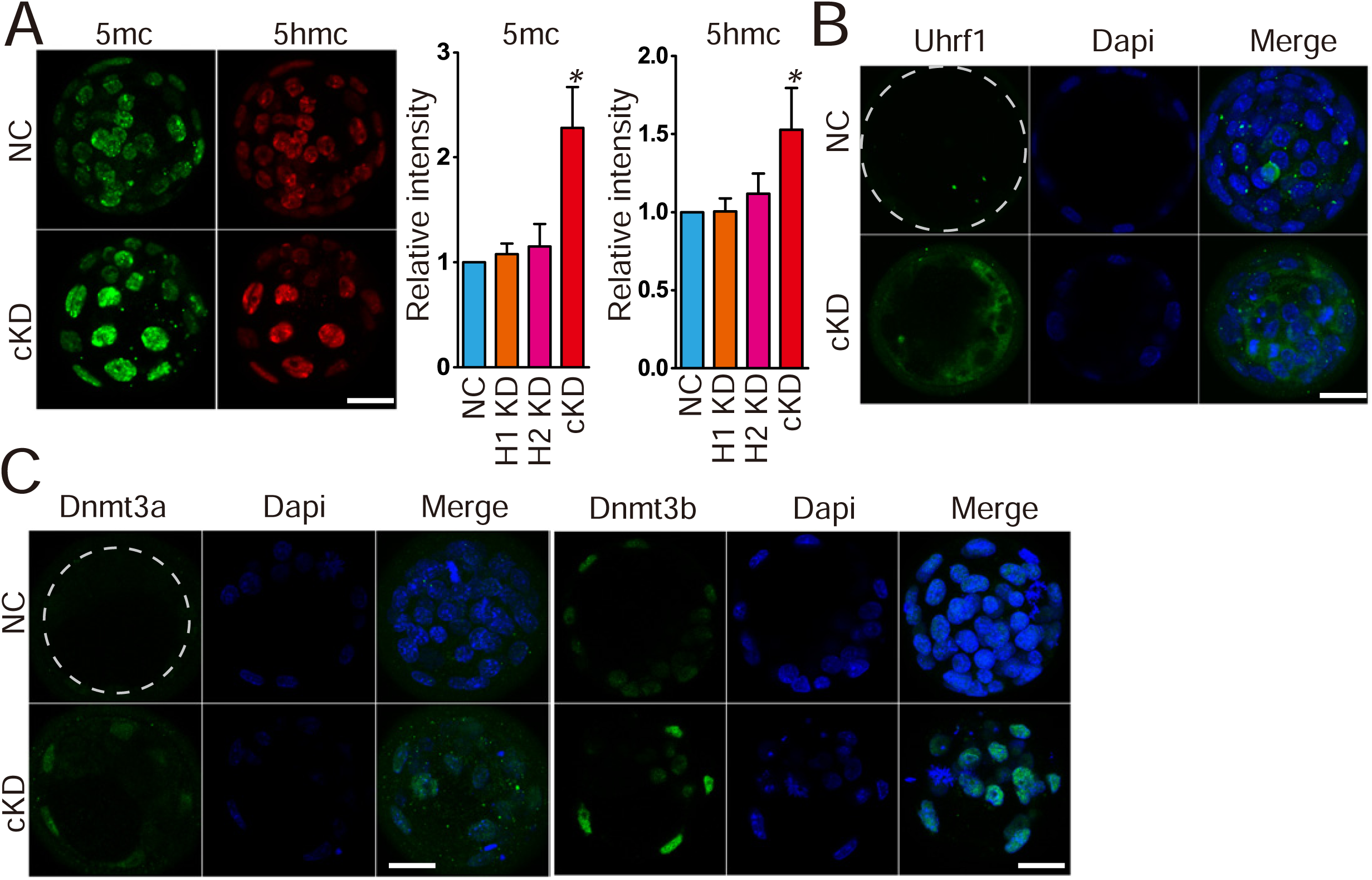
Increased global DNA methylation with upregulated DNA methyltransferases in embryos deficient of Hdac1 and 2. (A) Immunocytochemical analysis of 5’ methylcytosine (5mc) and 5 hydroxmethylcytosine (5hmc) in blastocysts. Both the intensity of 5mc and 5hmc was increased in cKD embryos (C and E) (n=3; 5-10 embryos were analyzed per group each time,, *P<0.05). (B-C) Hdac1 and 2 deficiency results in increased intensity of Uhrf1, Dnmt3a and Dnmt3b. The experiment was conducted three times and 8-10 embryos analyzed per group (*P<0.05; Scale bar: 25 µm).

Previous studies report Hdac1 physically interacts with DNA methyltransferases (Dnmts) and regulates the stability of Dnmt1 (35–38). There are three Dnmts present in preimplantation embryos: Dnmt1, 3a and 3b. Uhrf1 is a Dnmt1-interacting protein involved in the recruitment of Dnmt1 to maintain DNA methylation (39). The amount of Uhrf1 was increased not only in the nuclear but also in the cytoplasm in cKD blastocysts relative to control (n=3; Fig 7*B*). We have not found Dnmt1 antibody available for IF. However, immunoblotting analysis revealed an increase in Dnmt1 abundance when Hdac1/2 were inhibited (Fig S10*E*). In addition, Dnmt3a and 3b were barely detected in control mouse blastocysts whereas their signal intensity was significantly improved in cKD or FK228-treated embryos (Fig 7*C*, 7*D* and S10*D*). In summary, we conclude Hdac1 and 2 are critical for maintaining global DNA methylation properly through modulating the amount of Dnmts in preimplantation embryos.

### Double knockdown of Rbbp4 and 7 results in similar phenotypes as Hdac1/2 cKD embryos

Rbbp4 and 7 (also known as RbAp48 and 46) are two homologous chromatin-binding proteins that interact with Hdac1/2 to form the core components of multiple transcriptional corepressors, including Sin3a, NuRD, and CoREST (15)(Fig 8*A*). Both proteins have direct interactions with histone tails and are potentially responsible for recruitment of Hdac1/2-containing complexes to target sites. We next performed RNAi experiment to examine the functional consequences after knocking down Rbbp4 and 7. Effectiveness of siRNAs targeting *Rbbp4* and *7* was verified by IF analysis (Fig S11*A* and *B*). Analysis of embryogenesis *in vitro* showed that individual knockdown of Rbbp4 or 7 has no effect on preimplantation development, however, co-knockdown of Rbbp4 and 7 results in poor blastocyst rate and reduced total cell number per embryo at D4 (Fig 8*B* and *C*). The phenotype similarity between Rbbp4/7 cKD and Hdac1/2 cKD embryos prompted us to determine if defects in lineage specification and genome-wide methylation were also found. Both Cdx2 and Nanog were diminished in Rbbp4/7 cKD embryos (Fig 8*D* and *E*). An increase in global 5mc but not 5hmc was found in Rbbp4/7 cKD groups relative to controls (Fig 8*F*).

**Figure 8.**
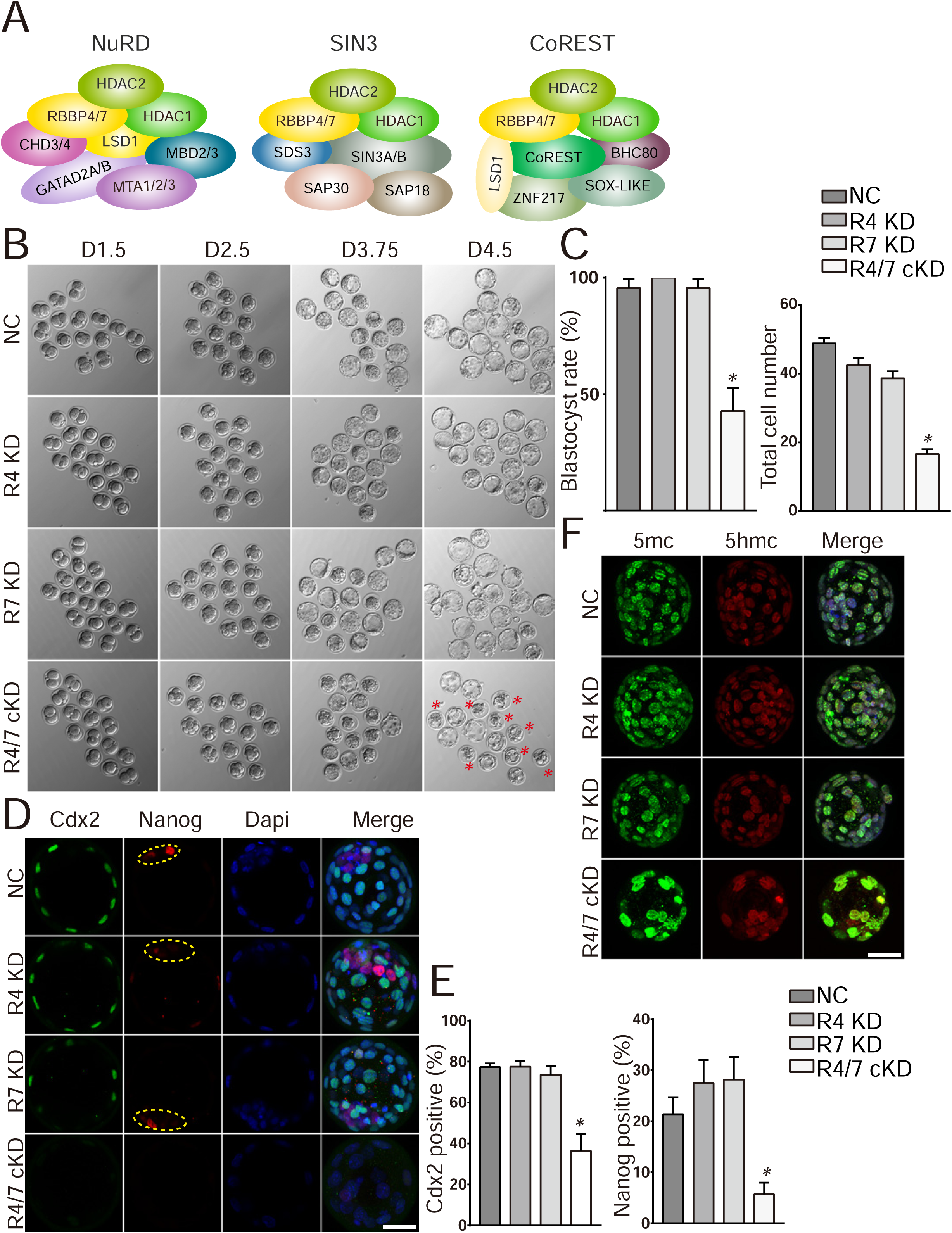
Rbbp4 and 7 deficiency leads to similar phenotypes as in Hdac1/2 cKD embryos. (A) Hdac1, Hdac2, Rbbp4, and Rbbp7 are core components in several epigenetic complexes: NuRD, Sin3, and CoREST. (B) Developmental potential of embryos lacking Rbbp4 and/or Rbbp7. Three replicates were conducted with 15-20 embryos analyzed per group per replicate. (C) Blastocyst rate and cell counting analysis of the experiment in panel B. (D) Both Cdx2 and Nanog were diminished in Rbbp4 and 7 cKD embryos. (E) Cdx2 or Nanog positive blastomeres were reduced in Rbbp4 and 7 cKD embryos. (F) Immunostaining analysis of 5mc and 5hmc.

## DISCUSSION

This report demonstrates that there is a functional redundancy for Hdac1 and Hdac2 in supporting preimplantation development. Depletion of both Hdac1 and 2 results in embryonic arrest during the morula to blastocyst transition with greatly disrupted transcriptome-wide expression profiles. Importantly, we document defects in three critical molecular events. First, Trp53 acetylation was induced and may contribute to increased apoptosis and cell cycle arrest. Second, lineage specification that generates TE and ICM was dramatically perturbed with defects including suppressed *Cdx2* expression and aberrant Hippo pathway. And third, a global increase of DNA methylation. Taken together, the combination of these effects contributes to the developmental failure of cKD embryos.

Double knockdown of Hdac1 and 2 in mouse preimplantation embryos results in developmental failure to pass blastocyst stage (Fig 1*E*). Previous studies and our present results indicate that independent knockdown of *Hdac1* or *Hdac2* does not affect blastocyst formation, suggesting a dispensable role during preimplantation development (22, 30). However, the developmental failure of cKD embryos suggests the viability of *Hdac1* or *Hdac2*-depleted embryos is due to functional redundancy of these closely related genes. In particular, we found compensatory roles of Hdac1 and Hdac2 in regulation of lineage specification, genome-wide methylation, and expression of critical genes, such as Cdx2 and Nanog. Overall, these two enzymes function redundantly during preimplantation development.

Transcriptome profiles were disturbed in embryos deficient of Hdac1 and 2. Hdac1/2 cannot bind to DNA directly. However, they can be tethered to DNA by many distinct transcription factors including YY1 (40), p130 (41), and Trp53 (42). Moreover, Hdac1/2 are recruited to DNA as components of multiprotein complexes, including Sin3a, NuRD, and the CoREST, which are well known for their transcriptional repressor activity. These facts could be the reason that the majority of DEGs are upregulated genes after double knockdown of Hdac1 and 2.

Lysine acetylation occurs not only to histones but various non-histone proteins, such as mitochondrial and cytosolic proteins (43). Hdac1/2 could also act as a “eraser” of these non-histone acetylation events (43). Trp53 is one of these non-histone proteins that is subject to acetylation. Trp53 plays a central role in a variety of biological processes including cell cycle arrest, DNA damage repair, apoptosis and metabolic changes (44). Our results clearly revealed the direct role of Hdac1 in acetylation of Trp53. Recently, Ma et al. also demonstrated that double knockout of Hdac1 and 2 leads to increased Trp53 acetylation. Overall, these results suggest it is a conserved mechanism on the direct regulation of Hdac1 and 2 on Trp53 acetylation (31).

Total cell counting and IF analysis suggest a cell cycle arrest in G2/M phase in cKD embryos. Previous studies demonstrated Hdac1 and 2 are associated with cell cycle progression across different cell types or tissues (20, 25, 45, 46). For instance, loss of Hdac1 and 2 in dividing cells results in a cell cycle block at G1 phase, which is partly attributed to the rise of the CDK-inhibitors, including p21 and p57 (21). However, we found no difference in p21 and p57 expression in cKD embryos, suggesting a different cell cycle block mechanism.

During the morula to blastocyst transition, the first lineage specification occurs with the regulation of contractility and critical signaling pathways, including Hippo and Notch (33). Core lineage-specific transcription factors, including *Cdx2* (TE-specific), and *Oct4* and *Nanog* (ICM-specific), are initially stochastically expressed and are gradually confined to specific lineages. However, it remains poorly understood how the expression of these factors themselves is controlled. Deficiency of Hdac1 and 2 results in failure of both the TE and ICM differentiation program. A dramatic reduction in blastocyst rate was observed and those cKD blastocyst that do form fail to outgrow, suggesting a lacking of functional ICM and TE. At a molecular level, expression of key marker genes, *Cdx2*, *Nanog* and *Oct4* were suppressed. In particular, Nanog and Cdx2 signal was barely seen in cKD embryos. Although the intensity of Oct4 was normal, its localization was not restricted to a subset of cells. These results collectively suggest Hdac1 and 2 are master regulators of the first lineage specification. Both RNA-seq and qPCR results suggest Hdac1 and 2 are involved the transcription of these key lineage-specific genes (Fig 2*C* and 4*A*-*E*). Indeed, ChIP-seq analysis shows that Hdac1 is enriched in active genes in ES and TS cells, such as TE-specific genes *Cdx2*, *Elf5* and *Eomes* and pluripotency network genes *Oct4*, *Nanog* and *Sox2* (18). These results may warrant further investigation through low input Chip-seq to determine if Hdac1 and Hdac2 colocalize at these critical genes during preimplantation stages (47).

The Hippo signaling pathway plays a crucial role in the first lineage specification, in particular for TE-specific program (33). Loss of Tead4 leads to lethality with a failure to generate functional TE and triggers downregulation of Cdx2 (48). As a central component of Hippo pathway, Yap could switch between nucleus and cytoplasm, which is phosphorylation-dependent. Yap acts as a transcriptional activator of Tead4 to induce TE-specific genes (34). Both Tead4 and Yap are disrupted in cKD embryos. Lats1 and Lats2 are both upstream regulators of Yap1 (34). Their expression was also reduced in cKD embryos. Interestingly, RNA-seq results also displayed dysregulation of genes in the Hippo pathway (Fig 2*E*). Thus, the downregulation of Cdx2 in cKD embryos may be partly due to aberrant Tead4 and Yap1 expression.

Our results suggest Hdac1 and 2 are critical for maintaining correct DNA methylation pattern during preimplantation development. Genome-wide removal of DNA methylation (5mc) occurs during preimplantation development, contrasting with stable DNA methylation pattern in somatic cells. There are two types of DNA demethylation: passive DNA demethylation that is DNA-replication-dependent and active demethylation that is achieved by enzymatically driven reactions. Our results show that Hdac1 and 2 affect the global active demethylation through regulating Dnmts and Uhrf1, a critical protein for recruiting Dnmt1 to specific DNA sequences. Increased Dnmt3a, 3b, and Uhrf1 protein abundance could be explained by increased transcripts in cKD embryos. However, we cannot rule out the possibility that loss of Hdac1 and 2 may affect the stability of Dnmts in preimplantation embryos. In contrast, Ma et al. found conditional knockout of Hdac1 and 2 in mouse oocytes resulted in a global decrease in DNA methylation and particularly reduced nuclear associated Dnmt3a (38). The discrepancy suggests a developmental context-dependent role of Hdac1/2 in regulation of Dnmts.

Functional analysis of Rbbp4 and 7 suggest these two proteins are critical components for ensuring functionality of Hdac1/2-containing chromatin complexes. Hdac1/2 and Rbbp4/7 are shared among several critical transcriptional corepressor, including Sin3a, NuRD and CoREST (15). Rbbp4/7 interacts directly with histone tails (H3 and H4) and are promising candidates for recruiting these epigenetic complexes. Interestingly, our results indicate Rbbp4 and 7 play a redundant function essential for preimplantation development, similar with Hdac1/2 cKD. We propose that Hdac1/2-Rbbp4/7-containing complexes are critically required for preimplantation development. Indeed, our previous studies documented an essential role of Suds3, a component of Sin3a complex, during preimplantation development with critical functions in lineage specification (30).

In summary, we documented a compensatory role of Hdac1 and 2 during preimplantation development. Hdac1 and 2 are essential for the regulation of cell cycle progression and apoptosis, which is probably mediated through acetylation of Trp53. Additionally, Hdac1 and 2 are required for the first cell differentiation program with a critical role in controlling expression of key TE and pluripotency-specific genes. In the context of chromatin regulation, Hdac1/2 are involved in maintaining proper genome-wide DNA methylation with dramatic effects on the protein abundance of Dnmts. Last but not least, deletion of Rbbp4 and 7, both structural partners of Hdac1/2 in several epigenetic complexes, results in similar phenotypes as Hdac1/2 cKD. Further understanding the mechanisms of epigenetic control during these early key molecular events will help the development of tools to reduce early embryonic lethality and improve the success of reproductive cloning in mammals.

## MATERIALS AND METHODS

### Ethics Statement

All experiments involving lab animals were conducted according to the guidelines for the care and use of lab animals and approved by Zhejiang University.

### Mouse embryo culture

Superovulation in B6D2F1 (C57BL/6 × DBA2, Charles River) female mice (8-10 weeks old) was performed by injecting 10 IU PMSG (San-Sheng pharmaceutical Co. Ltd., Ningbo, China) followed by 10 IU hCG (San-Sheng pharmaceutical Co. Ltd., Ningbo, China) 46-48 h later. At 20-22 h post-hCG treatment, zygotes were collected from B6D2 F1 female mice mated to B6D2 F1 males. Hyaluronidase (Sigma, St Louis, MO, USA) was used to remove cumulus cells. Zygotes were cultured in KSOM at 37°C/5% CO_2_. For FK228 treatment experiment, mouse morula were treated with Romidepsin (FK228, Depsipeptide, Selleck, 50 nM) for 12 hours. For embryo transfer experiment, 2-cell stage control or cKD embryos were transferred into the oviduct of pseudo-pregnant female mice (ICR).

### Outgrowth

For outgrowth formation experiment, individual blastocyst was collected on D4, removed of zona pellucida or kept intact, and incubated in DMEM (Gibco) containing 10% FBS (Gibco) on 48-well plates coated with 0.1% Gelatin (Gibco).

### Microinjection

siRNAs and mRNAs were microinjected into the cytoplasm of zygote using a Piezo-drill (Eppendorf, Germany) and Eppendorf transferman micromanipulators. siRNA (20 µM; GenePharma, Shanghai) and/or synthetic mRNA (500 ng/µl) were loaded into microinjection pipette and constant flow was adjusted to allow successful microinjection. Approximately 10 pl of siRNA and/or mRNA was delivered into the cytoplasm of zygotes or 2 cell blastomere by microinjection. Sense and antisense sequences of siRNAs used in the present study was listed in Table S3.

### *In vitro* RNA synthesis

Wildtype cDNA for *Hdac1*, *Hdac2* and *H2B-RFP* were cloned into T7-driven vectors. *Hdac1* mutants (H141A) were constructed as described previously (31). All sequences were confirmed by Sanger sequencing prior to use. To prepare mRNAs for microinjection, expression vectors were linearized and then were *in vitro* transcribed, capped and poly(A) tailed using T7 mMESSAGE mMACHINE Ultra Kit (Life Technologies, Grand Island, NY, USA) based on the manual. mRNA was recovered and purified by MEGAclear Kit (Life Technologies, Grand Island, NY, USA) and the integrity validated by electrophoresis.

### TUNEL

The embryos were washed in 0.1% PVP/PBS, fixed in 4% PFA for 10 min and permeabilized in PBS containing 0.5% Triton X-100 and 0.1% sodium citrate for 30 min. Then, the samples were incubated in a buffer solution of TDT 10X, CoCl_2_, 2mM dATP, 0.5 units/μl terminal deoxynucleotidyl transferase enzyme and 0.5 mM FITC-dUTP for 1 h at 37°C in humidity. DNA was stained with DAPI. Samples were mounted onto slides and imaged with confocal microscope system (Zeiss LSM780).

### Immunofluorescence

Preimplantation embryos were fixed with 4% paraformaldehyde in PBS for 10 min at room temperature, permeabilized with 0.5% Triton X-100 for 30 min, then blocked in 10% FBS/0.1% Triton X-100/PBS for 1 h after 3 times washing in 0.1% Triton X-100 PBS, and incubated with antibodies (Table S3) 1 h at room temperature or overnight at 4°C followed by incubation with Alexa Flour secondary antibodies 488, 595 (Invitrogen) at 37°C for 1 h. DNA was stained with DAPI and samples were mounted and observed with a Zeiss LSM780 confocal microscope (Zeiss).

### Reverse transcription and real time PCR

Total RNA from embryos was extracted using the Arcturus Picopure RNA isolation kit (Life Technologies, Grand Island, NY, USA). cDNA synthesis was performed using a reverse transcription system (Invitrogen). To quantify gene expression differences between KD and control groups, real-time PCR was performed on a StepOne^TM^ system using using FastStart Universal SYBR Green Master (Roche). *H2a* was used as an endogenous control.

### Western blotting

Embryos were lysed on ice in RIPA lysis buffer (Beyotime) supplemented with 1 mM phenylmethylsulfonyl fluoride (Beyotime). Equal numbers of embryos were used in each group. Protein were separated by 8% SDS-PAGE and transferred to a polyvinylidene fluoride membrane (Millipore). Then, membrane was blocked with 5% non-fat milk and incubated with primary antibodies overnight at 4℃ and secondary antibodies for 1.5 h at room temperature. Signals were detected with WESTAR NOVA 2.0 (Cyanagen).

### RNA-seq and bioinformatic analysis

At E2.75, embryos were collected from NC and cKD groups (60 embryos per sample, n=3). Total RNA was isolated from embryos using Picopure RNA isolation kit (Life Technologies, Grand Island, NY, USA) according to the manufacturer’s instruction. Before RNA extraction, 2 × 10^6^ copies of RFP and GFP mRNA was added. mRNAs were separated with oligo(dT)25 beads, and was used to prepare sequencing libraries with NEB Next Ultra RNA Library Prep Kit for Illumina (New England Biolabs). Briefly, mRNA was fragmented and reverse transcribed. The cDNA library was subject to end repair, poly(A)-tailing, adaptor ligation, and PCR amplification of 12–15 cycles for sequencing library construction. The library was sequenced by Illumina Hiseq X Ten and RNA-seq reads were assigned directly to transcripts and counted with Salmon (https://combine-lab.github.io/salmon/)(49, 50). Differential expression analysis was performed by DESeq2 package with P adjusted <0.05 and fold change >2 or <0.5). GO and KEGG analysis for enrichment of differentially expressed genes was determined using the Database for Annotation, Visualization and Integrated Discovery (DAVID).

### Statistical Analysis

Differences between two groups were determined by two-tailed unpaired Student’s t tests. All experiments were repeated at least three times unless otherwise stated. For quantification of IF results, nuclear areas were outlined based on DAPI signal and mean intensity measured using NIH ImageJ. Signal intensities were normalized to control embryos. A value of P<0.05 was considered to be statistically significant. RNA-seq results were analyzed with R (http://www.rproject.org). Results are stated as mean ± S.E.M.

## Acknowledgments

We thank all members of the K. Zhang laboratories for their helpful discussions; S. Hong (Lab Animal Core Facility, Zhejiang University) for help in embryo transfer; Alan D. Ealy (Virginia Tech, Blacksburg, Virginia, USA) for critically reading the manuscript. This work was supported by National Natural Science Foundation of China (No. 31672416 and No. 31872348) and the Foundation of Key Laboratory of Veterinary Biotechnology (No. klab201708), Shanghai, China.

---

**Figure S1.**
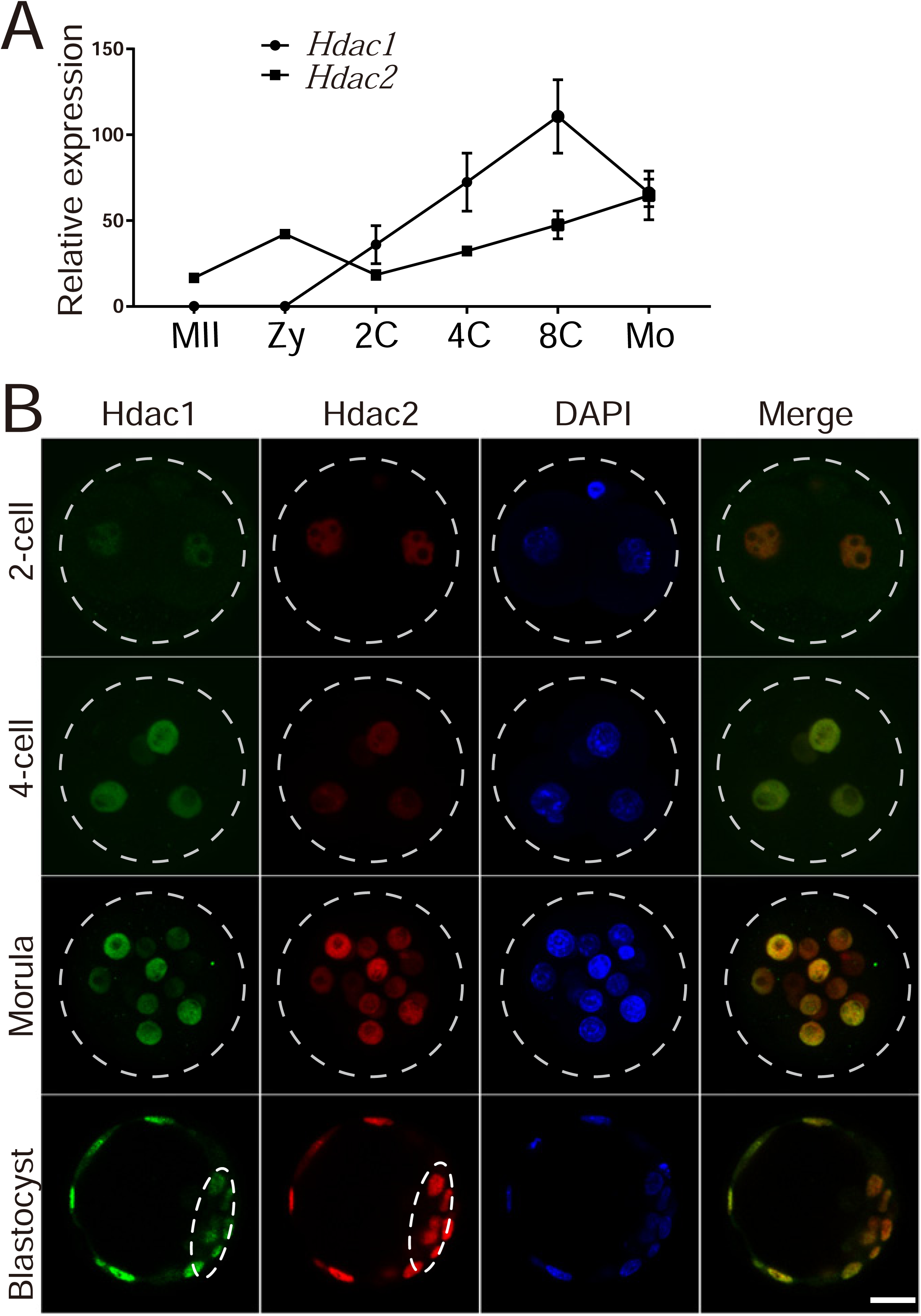
Extensive expression and colocalization of Hdac1 and 2 through preimplantation development. (A) Analysis of expression level of Hdac1 and 2 from a previous published single cell RNA-seq datasets (GSE44183). (B) Immunocytochemical detection of Hdac1 and 2 through preimplantation development. At least 10 embryos were analyzed and the experiment was performed three times. Scale bar: 25 µm.

**Figure S2.**
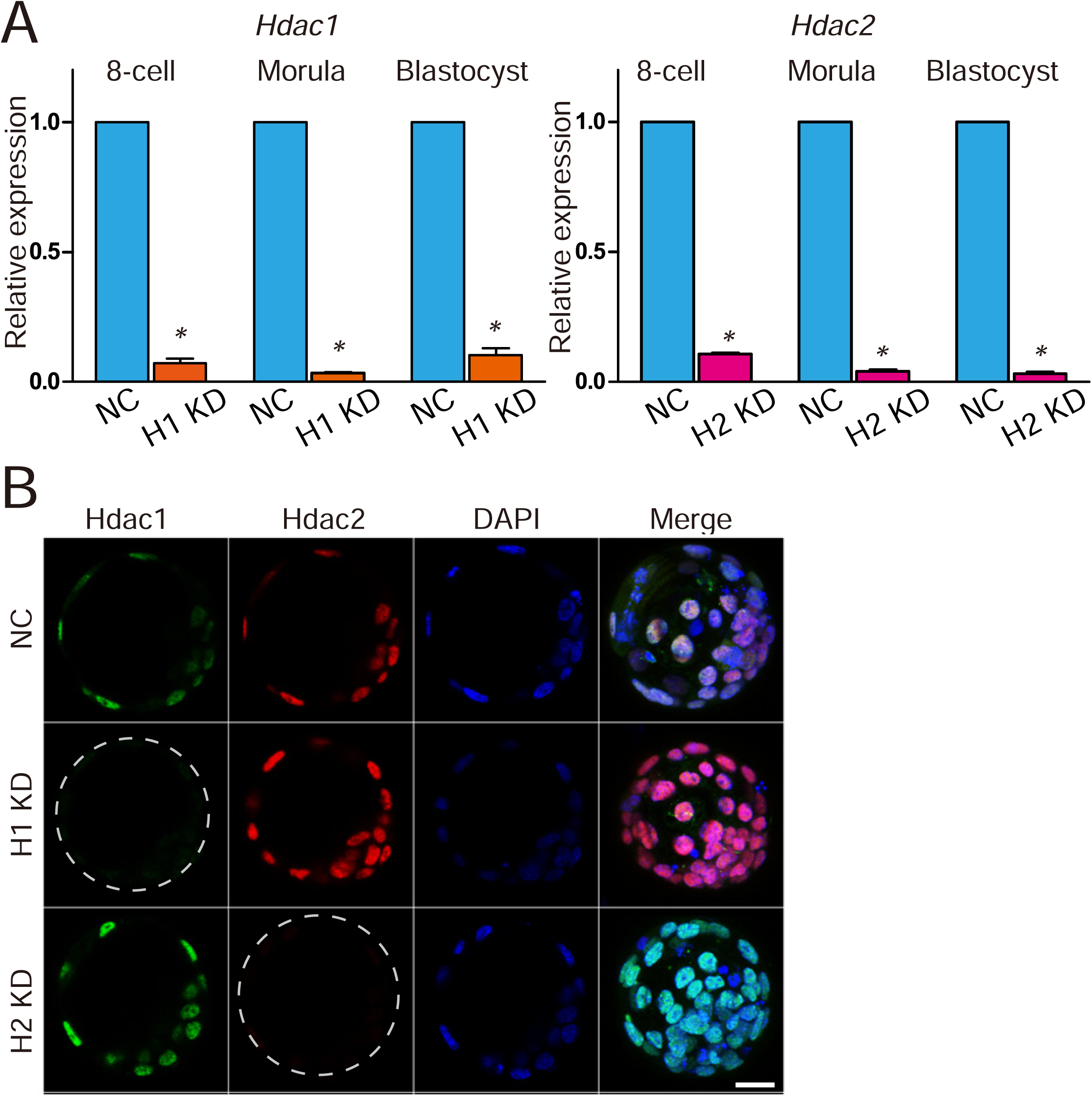
Validation of knockdown efficiency of siRNAs targeting *Hdac1* or *2*. (A) qPCR analysis of *Hdac1* or *2* in Hdac1 or Hdac2 KD embryos, respectively (n=3 pools of 5-10 embryos each per group, *P<0.05). (B) Immunocytochemical analysis of Hdac1 and 2 in Hdac1 or Hdac2 KD embryos (n=3; 5-10 embryos were analyzed per group each time, *P<0.05). Scale bar: 25 µm.

**Figure S3.**
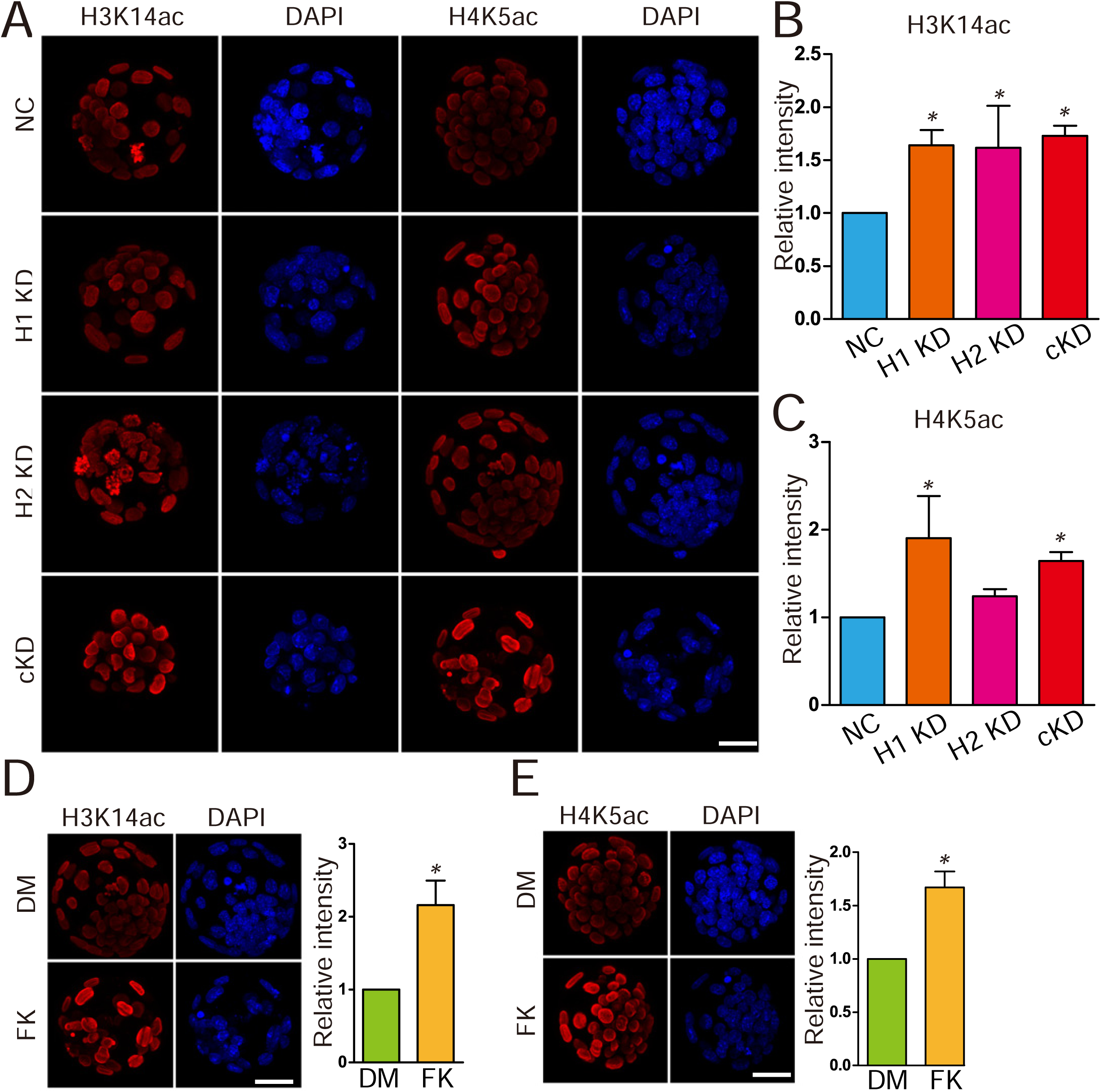
Hdac1/2 co-knockdown or FK228 treatment results in increased intensity of histone H3 lysine 14 acetylation and histone H4 lysine 5 acetylation. (A-E) Immunocytochemical analysis of histone H3 lysine 14 acetylation (H3K14ac) and histone H4 lysine 5 acetylation (H4K5ac) in embryos after RNAi (A-C) or FK228 treatment (D and E) (6-10 embryos were analyzed per group, *P<0.05). Scale bar: 25 µm.

**Figure S4.**
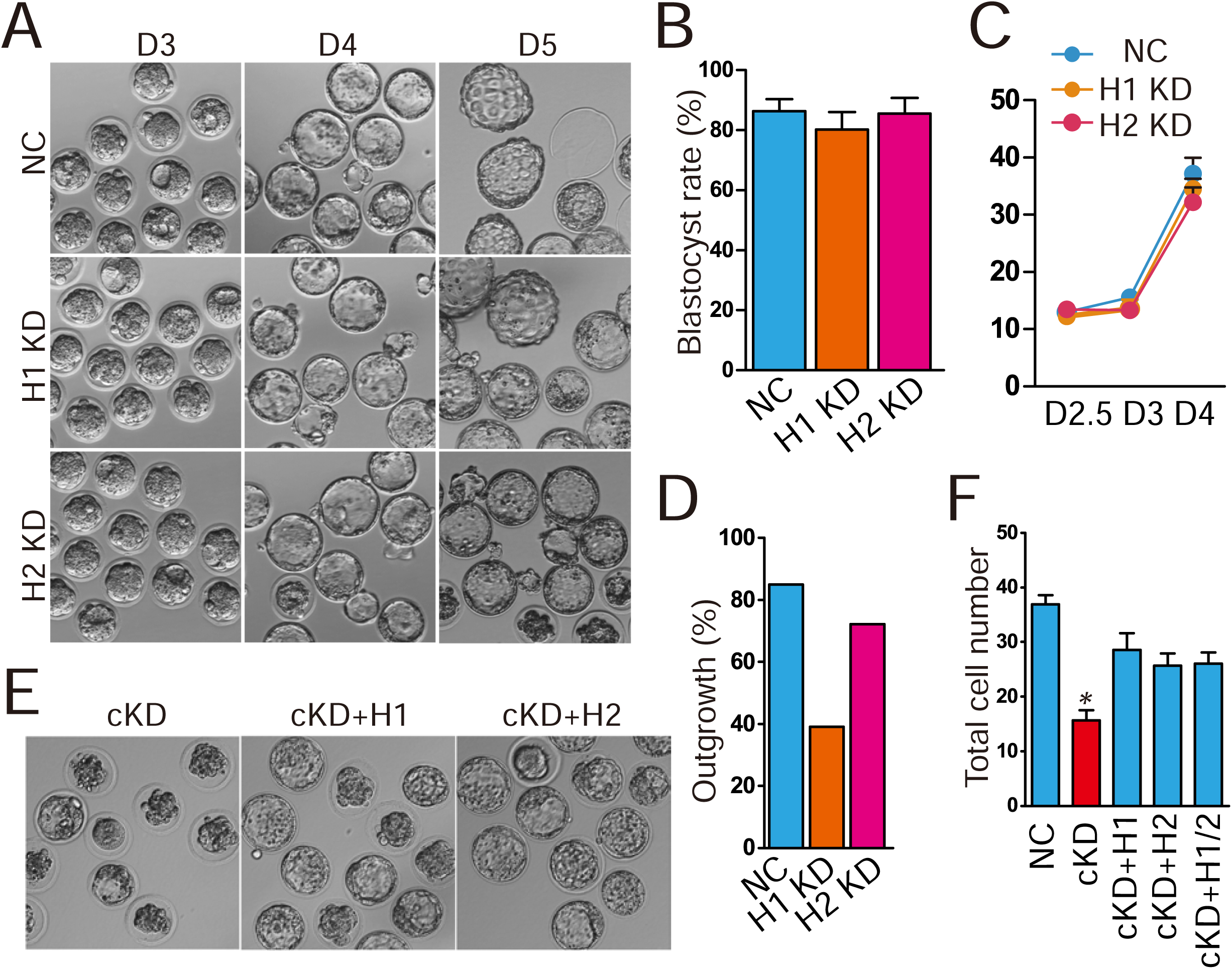
Effects of Hdac1 or 2 individual KD on preimplantation development. (A) Representative photos of NC, Hdac1 (H1) KD, and Hdac2 (H2) KD embryos from D3 to D5. No difference was noted between H1 or H2 KD and NC group through preimplantation development. (B and C) Blastocyst rate (B) at D4 and total cell number per embryo (C) was similar among NC, H1 KD, and H2 KD groups (n=3; 15-20 embryos analyzed per group). (D) Incidence of outgrowth formation for blastocysts derived from NC, H1 KD, and H2 KD embryos (20-25 embryos analyzed per group). (E and F) Rescue of cKD embryos by injection of exogenous Hdac1 or Hdac2 mRNA (n=3).

**Figure S5.**
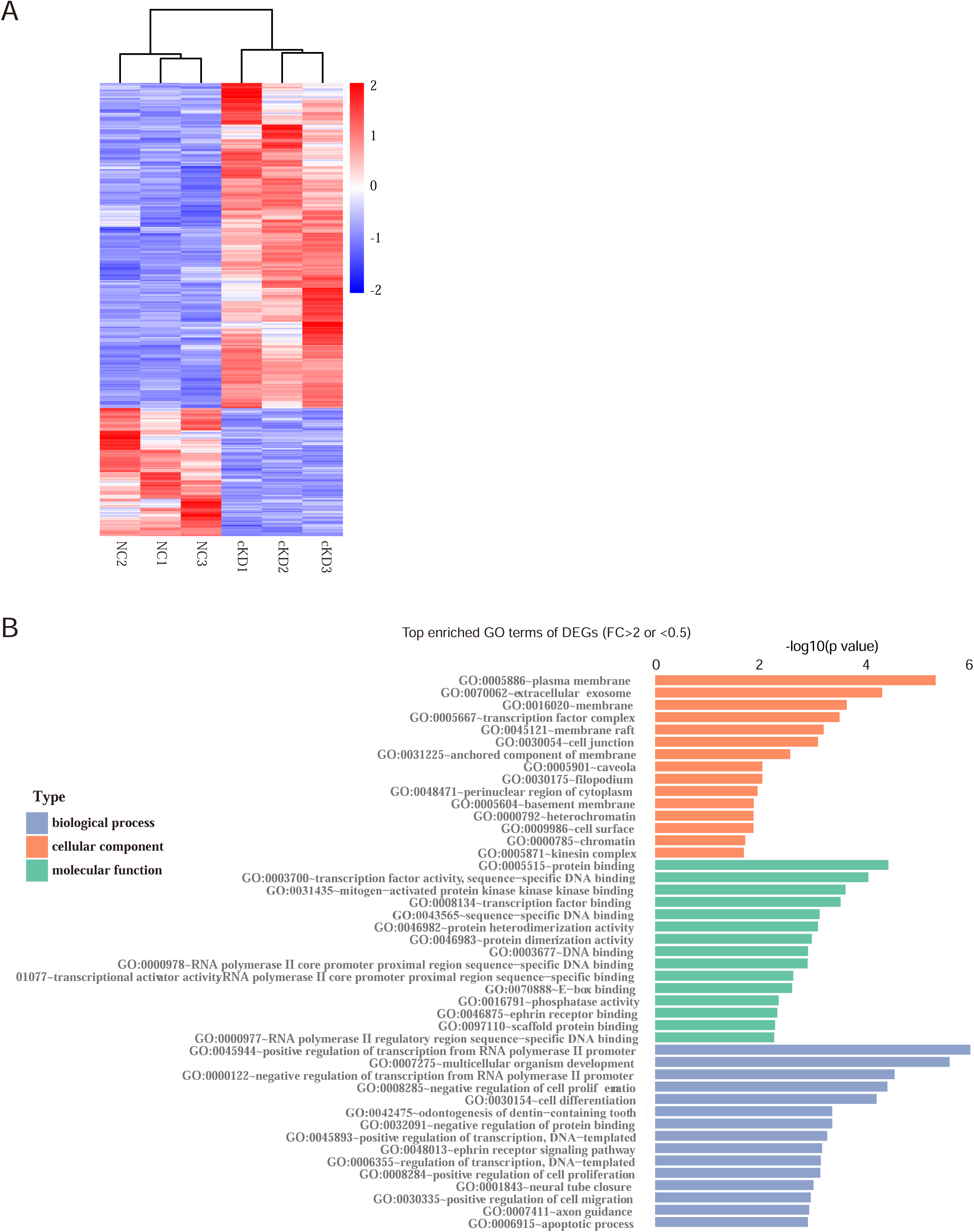
Transcriptomic analysis of embryos deficient of Hdac1 and 2. (A) Heatmap showing differentially expressed genes (DEG) between cKD and NC embryos. (B) GO analysis of all DEGs in cKD embryos related to NC groups.

**Figure S6.**
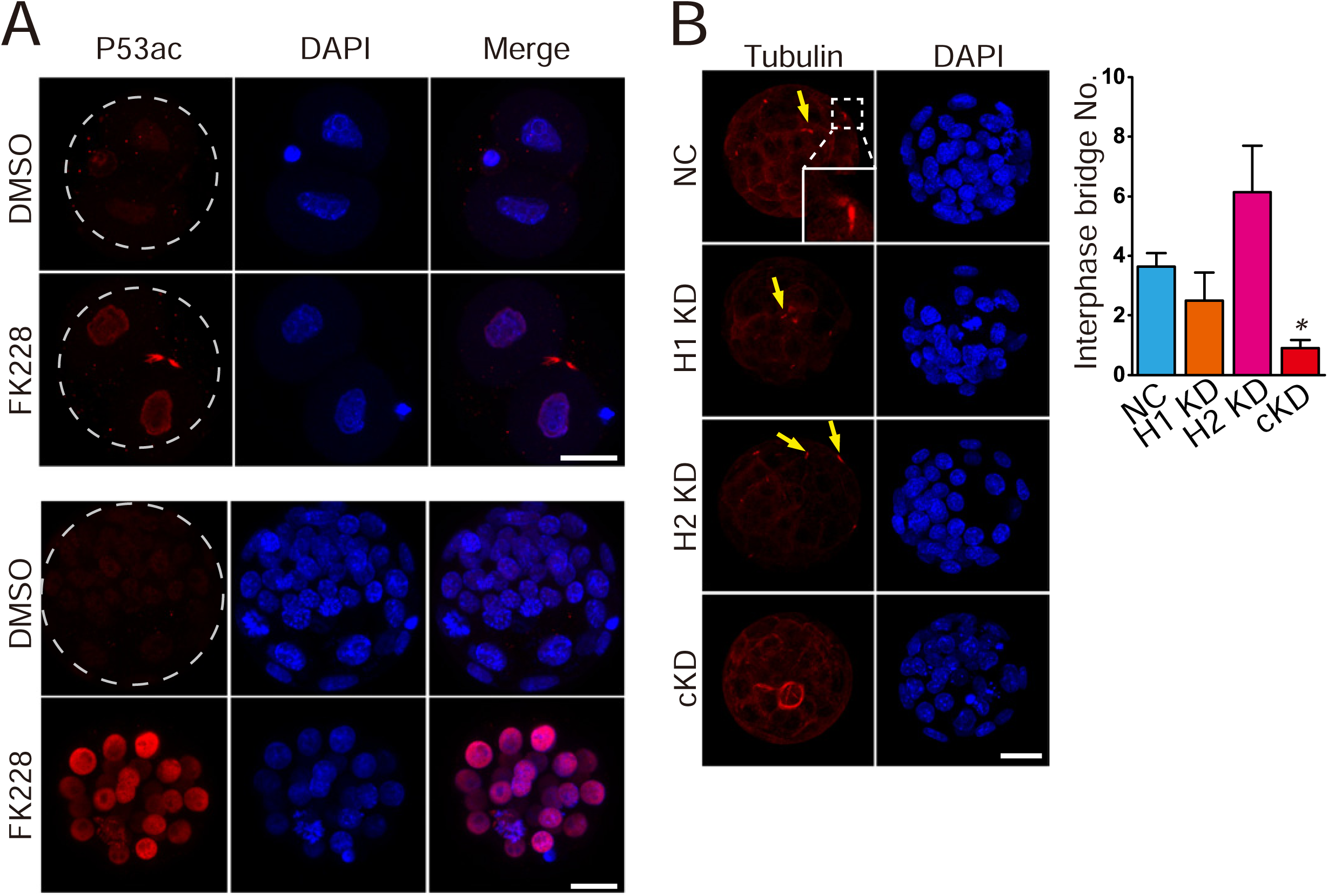
Blockage of Hdac1 and 2 results in increased Trp53 acetylation and aberrant interphase bridge. (A) Immunocytochemical analysis of Trp53K379 acetylation (P53ac) in 2-cell embryos and morula. Mouse zygotes or morula were treated with FK228 for 12 h. The intensity of P53ac was increased dramatically in FK228-treated embryos. (E) Immunocytochemical examination of α-tubulin, a marker for interphase bridge, in blastocysts (Arrows: interphase bridge; n=3; 5-10 embryos were analyzed per group each time, Scale bar: 25 μm).

**Figure S7.**
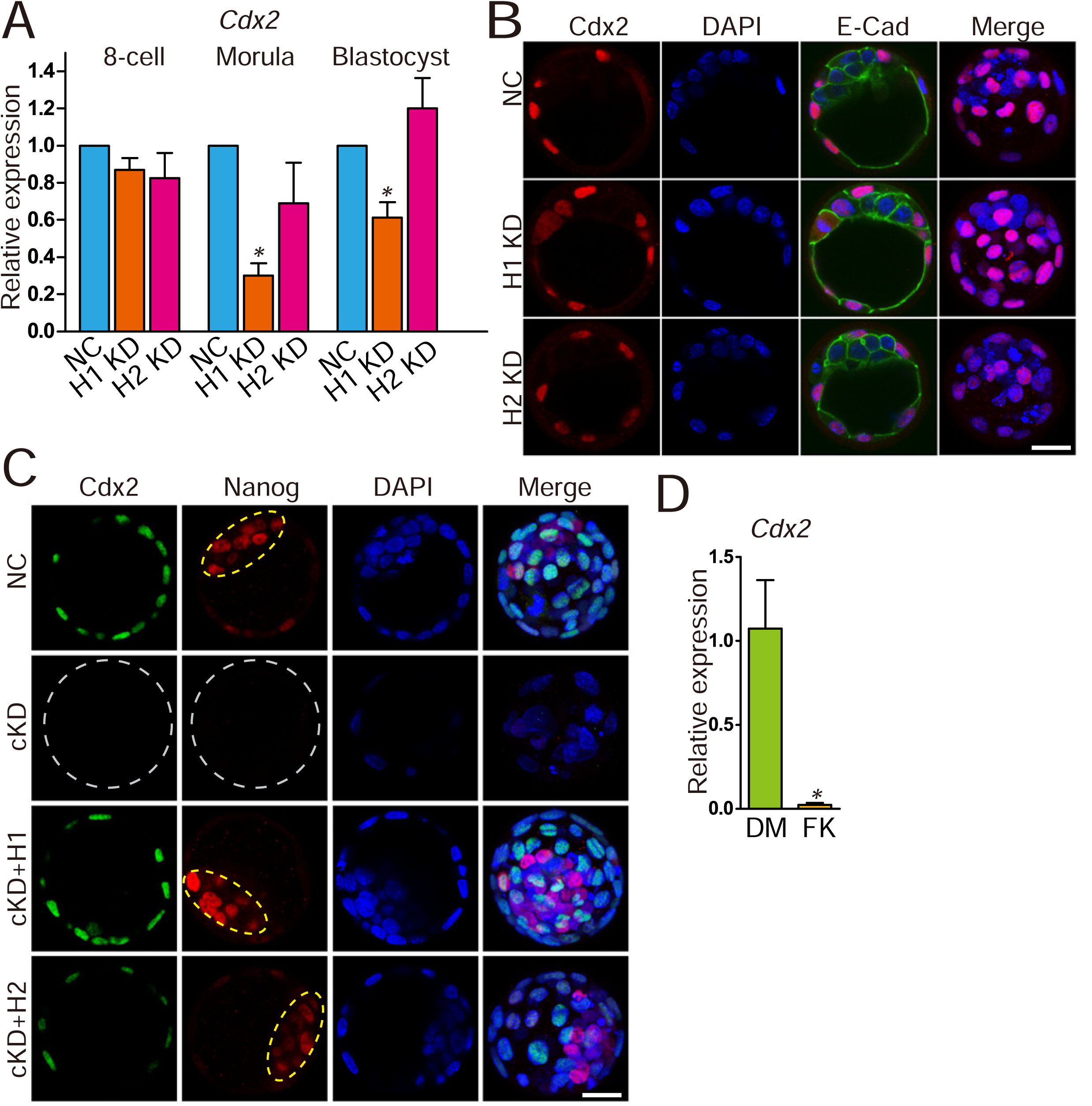
Effects of Hdac1 KD or Hdac2 individual KD on Cdx2 expression. (A) qPCR analysis of *Cdx2* in 8-cell embryos, morula and blastocysts (n=3 pools of 5-10 embryos each per group; *P<0.05). (B) Immunocytochemical analysis of Cdx2 in morula and blastocysts. The intensity of Cdx2 was not changed in H1 or H2 KD embryos (n=3; 5-10 embryos were analyzed per group each time). (C) Rescue of Cdx2 in cKD embryos after injection of exogenous *Hdac1* and/or *2*. The experiment was conducted three times and 5-10 embryos analyzed per group per time. (D) FK228 treatment results in significant decrease in Cdx2 expression level (n=3; 5-10 embryos were analyzed per group each time, *P<0.05). Scale bar: 25 μm.

**Figure S8.**
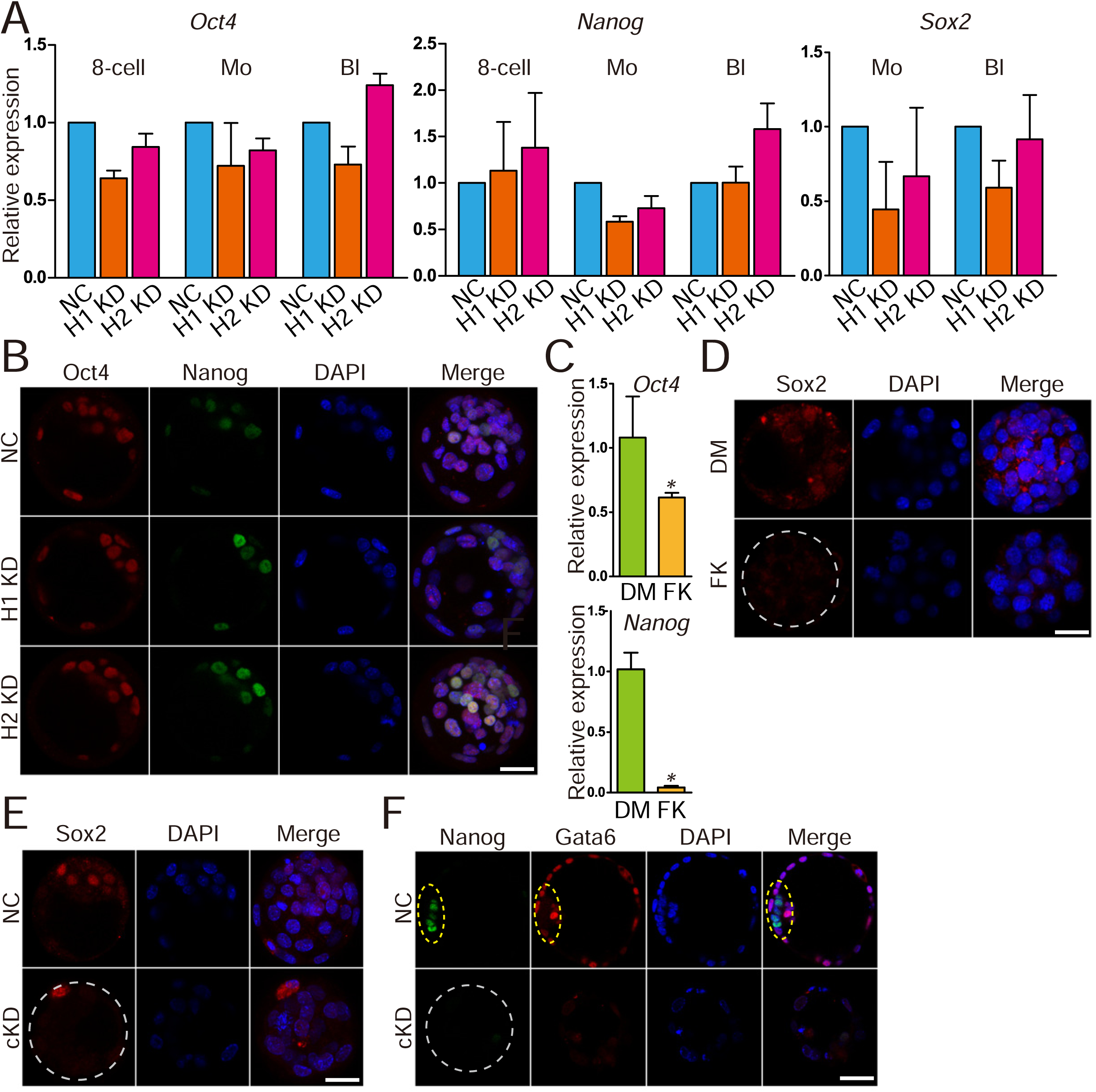
Effects of Hdac1 KD or Hdac2 individual KD on Oct4, Nanog and Sox2. (A) qPCR analysis of *Oct4, Nanog and Sox2* in NC, Hdac1 KD, and Hdac2 KD embryos. (B) Immunocytochemical analysis of Oct4 and Nanog in blastocysts. (C) qPCR analysis of *Oct4* and *Nanog* in FK228-treated embryos. (D and E) Immunocytochemical analysis of Sox2 in cKD or FK228 treated embryos. (F) Immunostaining detection of Nanog and Gata6 in late blastocysts.

**Figure S9.**
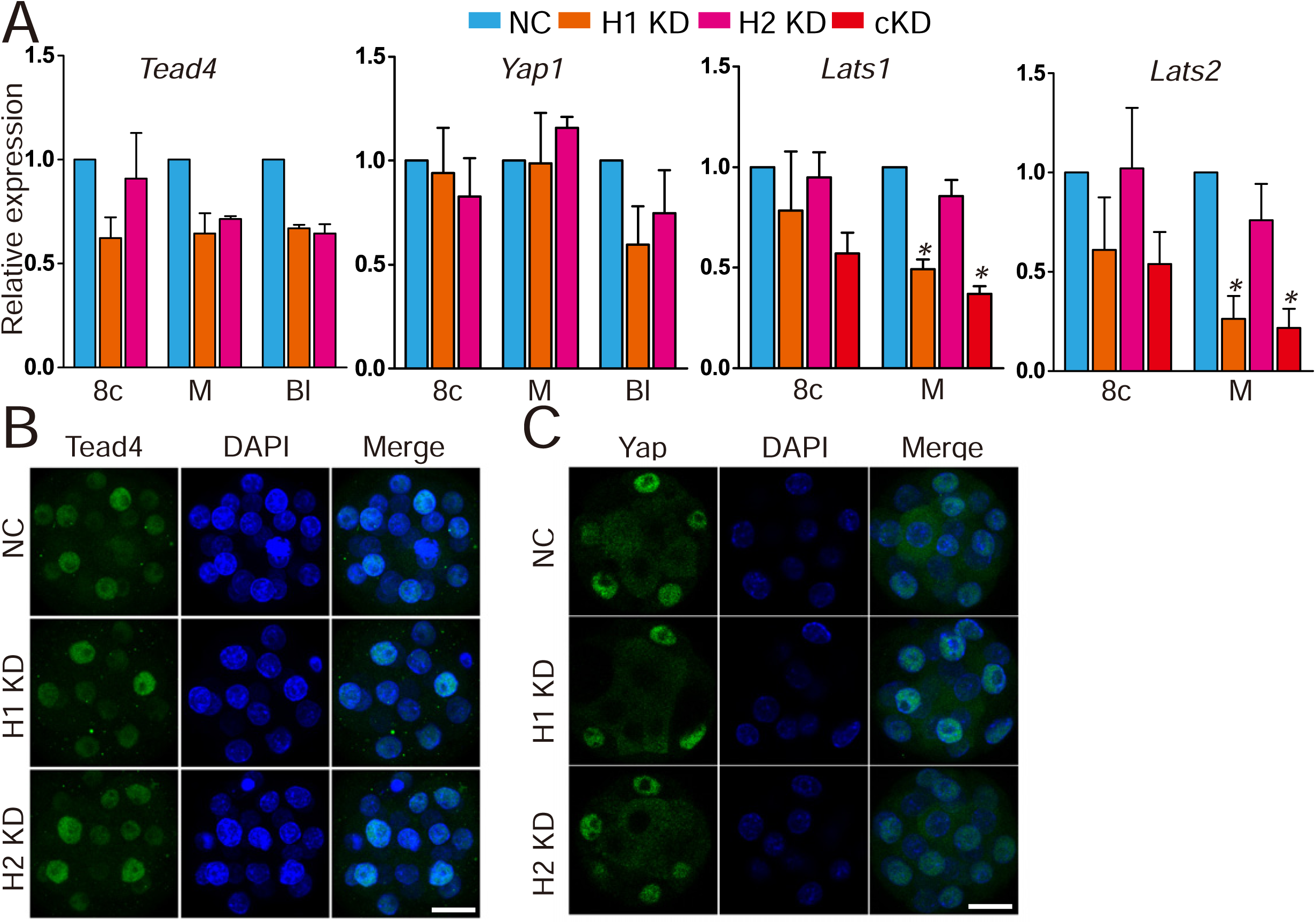
Effects of Hdac1/2 individual KD on Hippo signaling pathway. (A) qPCR analysis of *Tead4*, *Yap1, Lats1,* and *Lats2* in NC, Hdac1 KD, and Hdac2 KD embryos. (B and C) Immunocytochemical analysis of Tead4 and Yap in morula deficient of Hdac1 or Hdac2.

**Figure S10.**
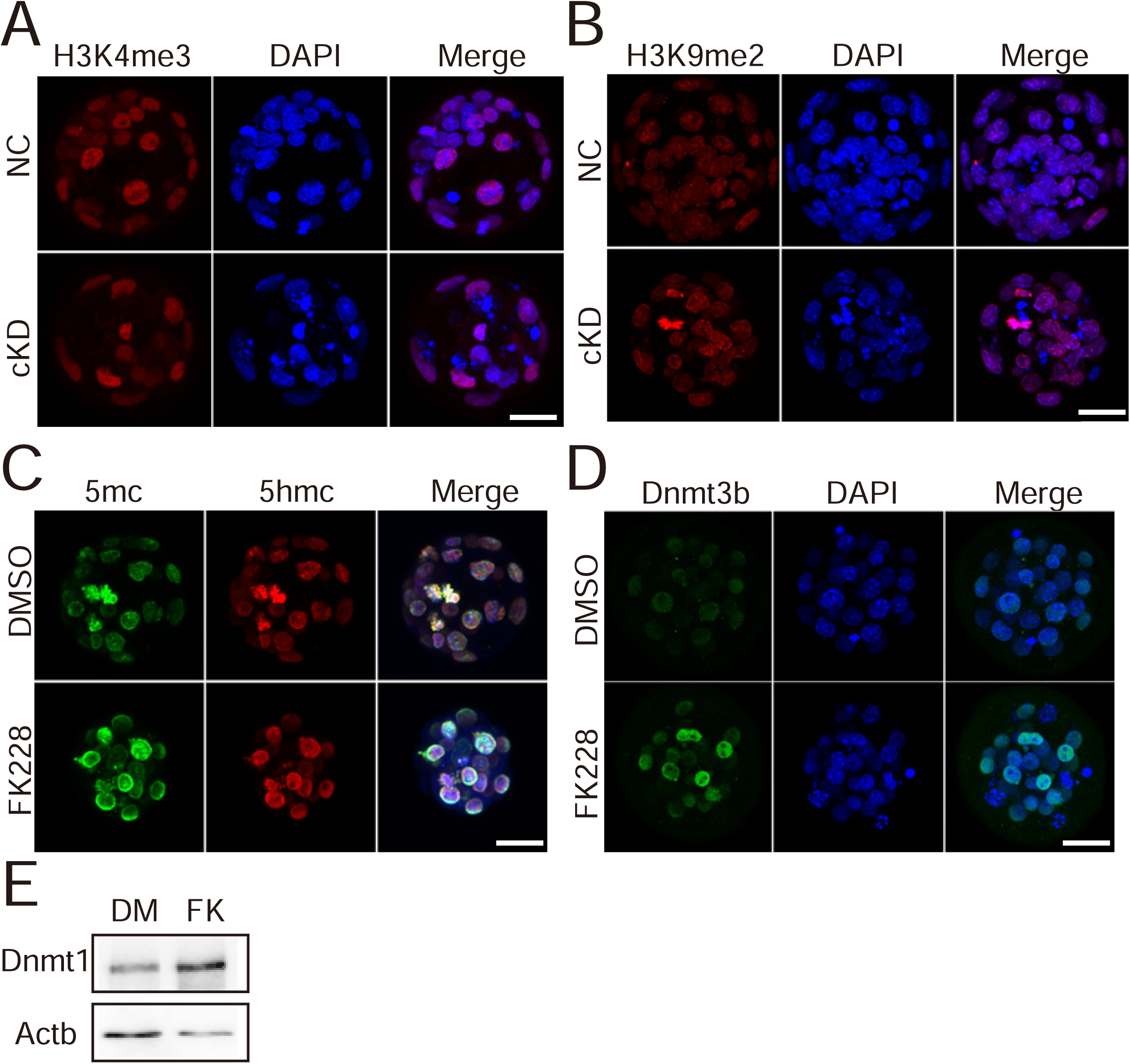
Effects of Hdac1 and 2 cKD on epigenetic modifications. (A and B) Immunocytochemical analysis of H3K4me3 and H3K9me2 in morula deficient of Hdac1 and Hdac2. (C and D) FK treatment results in an increase of DNA methylation (5mc) and Dnmt3b intensity. (E) Immunoblotting analysis of Dnmt1 in morula/blastocysts treated with DMSO (DM) or FK228 (FK). Two replicates were performed with 40 embryos used per group and similar results obtained.

**Figure S11.**
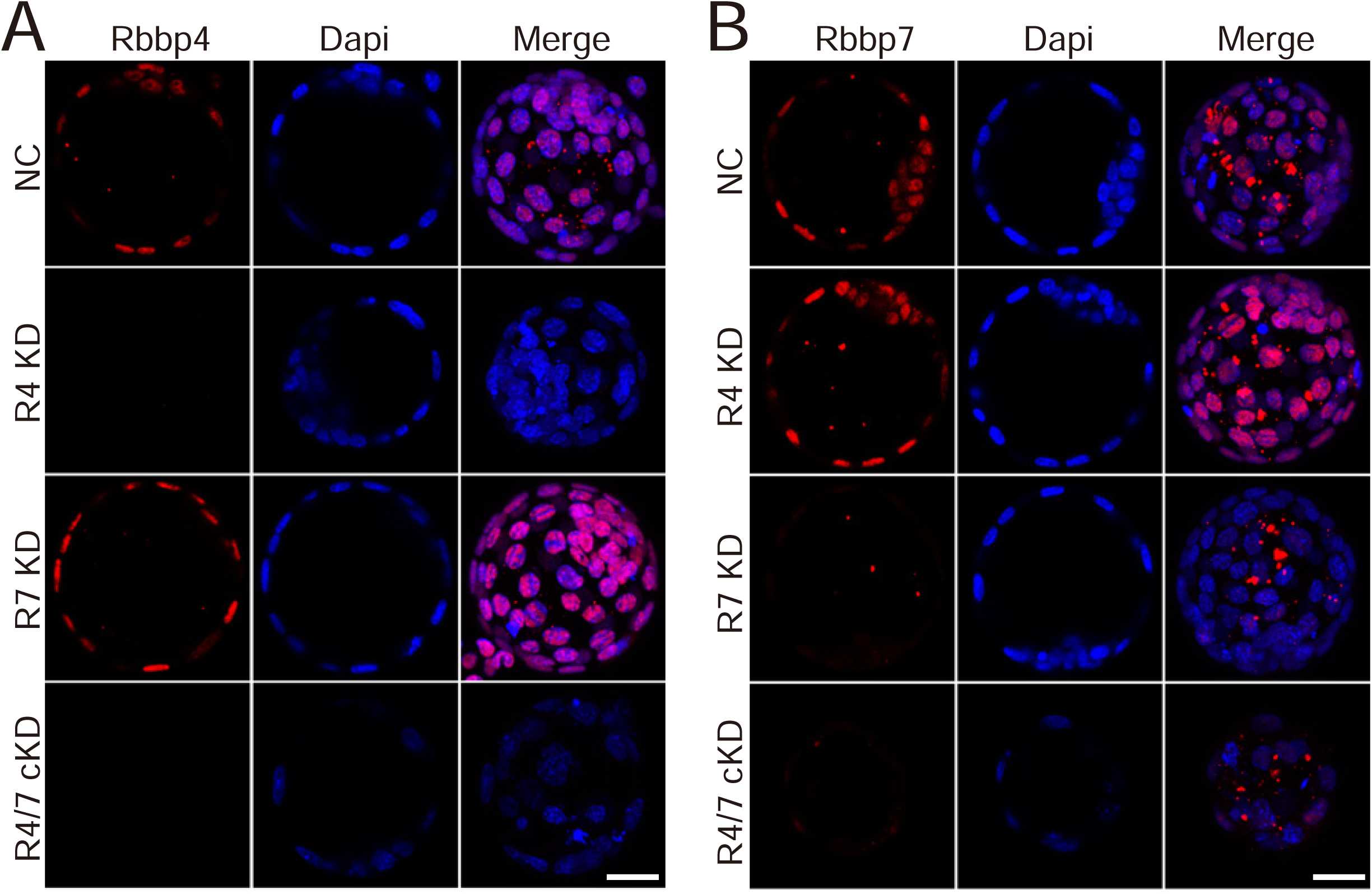
Validation of knockdown efficiency of siRNAs targeting *Rbbp4* or *7*. (A-B) Immunocytochemical analysis of Rbbp4 and 7 in Rbbp4 or 7 KD embryos. Scale bar: 25 µm.

**Table S1.** List of differentially expressed genes between control and Hdac1 and 2 cKD embryos (Supporting Excel file).

**Table S2.**
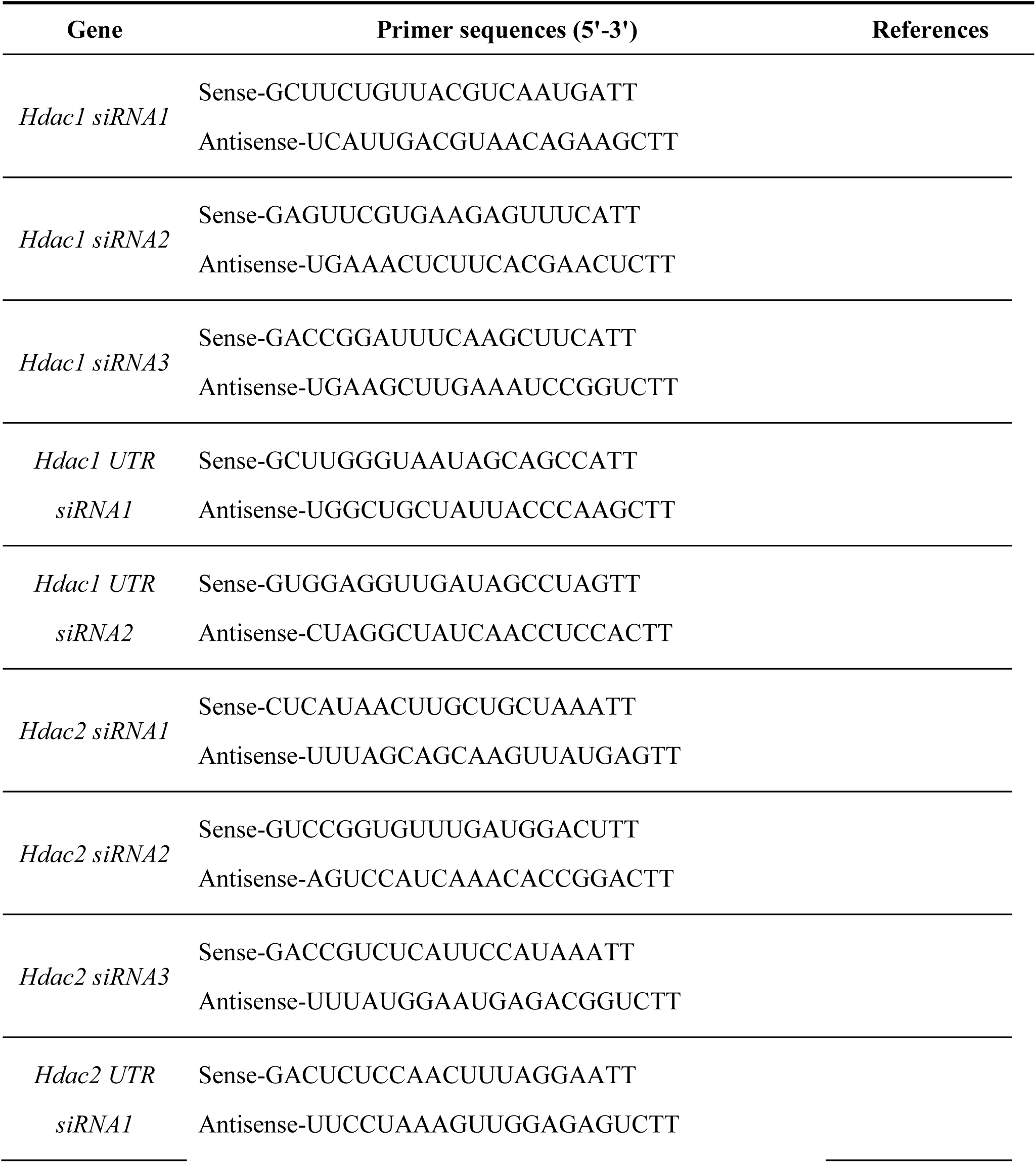
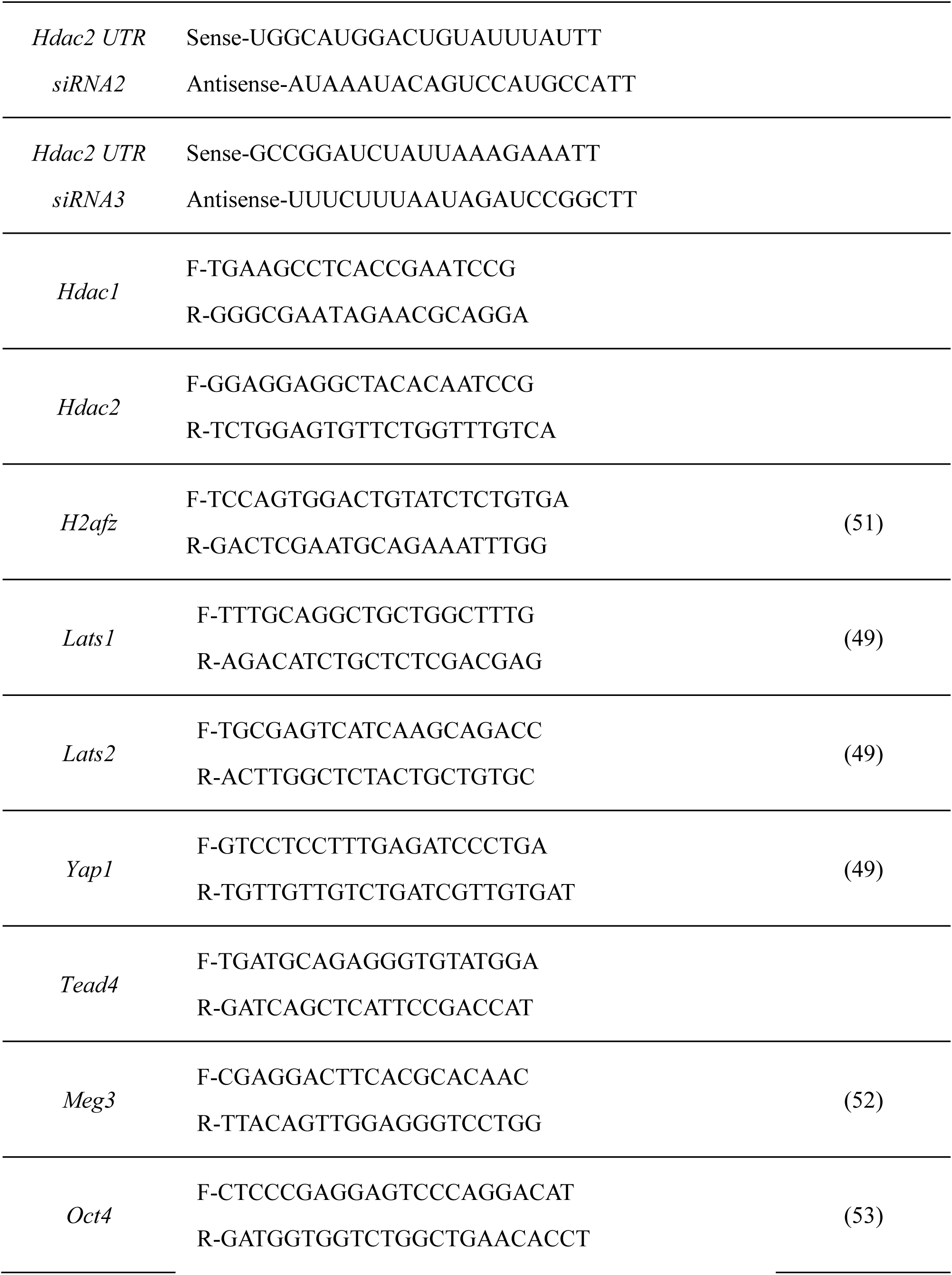
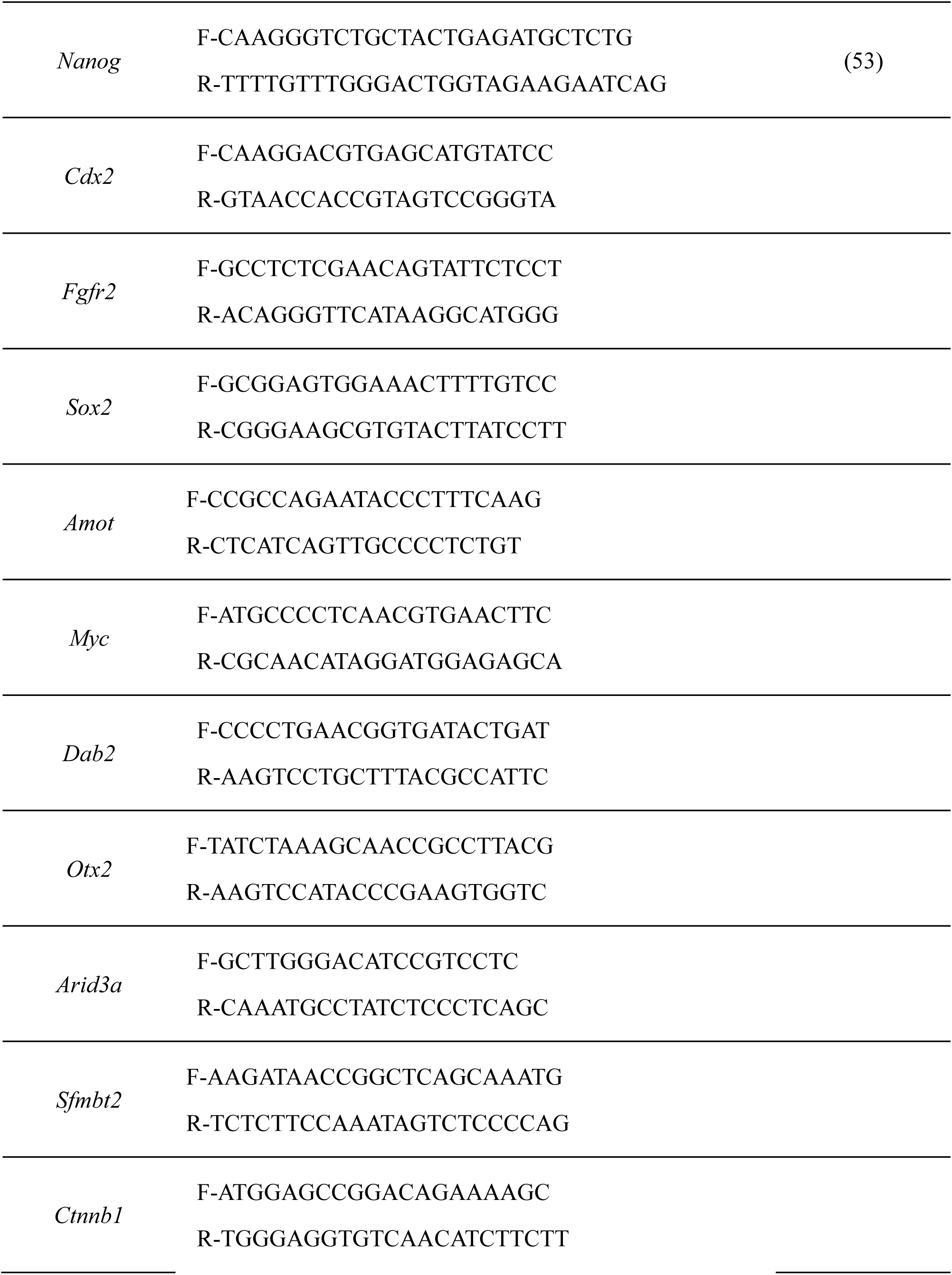
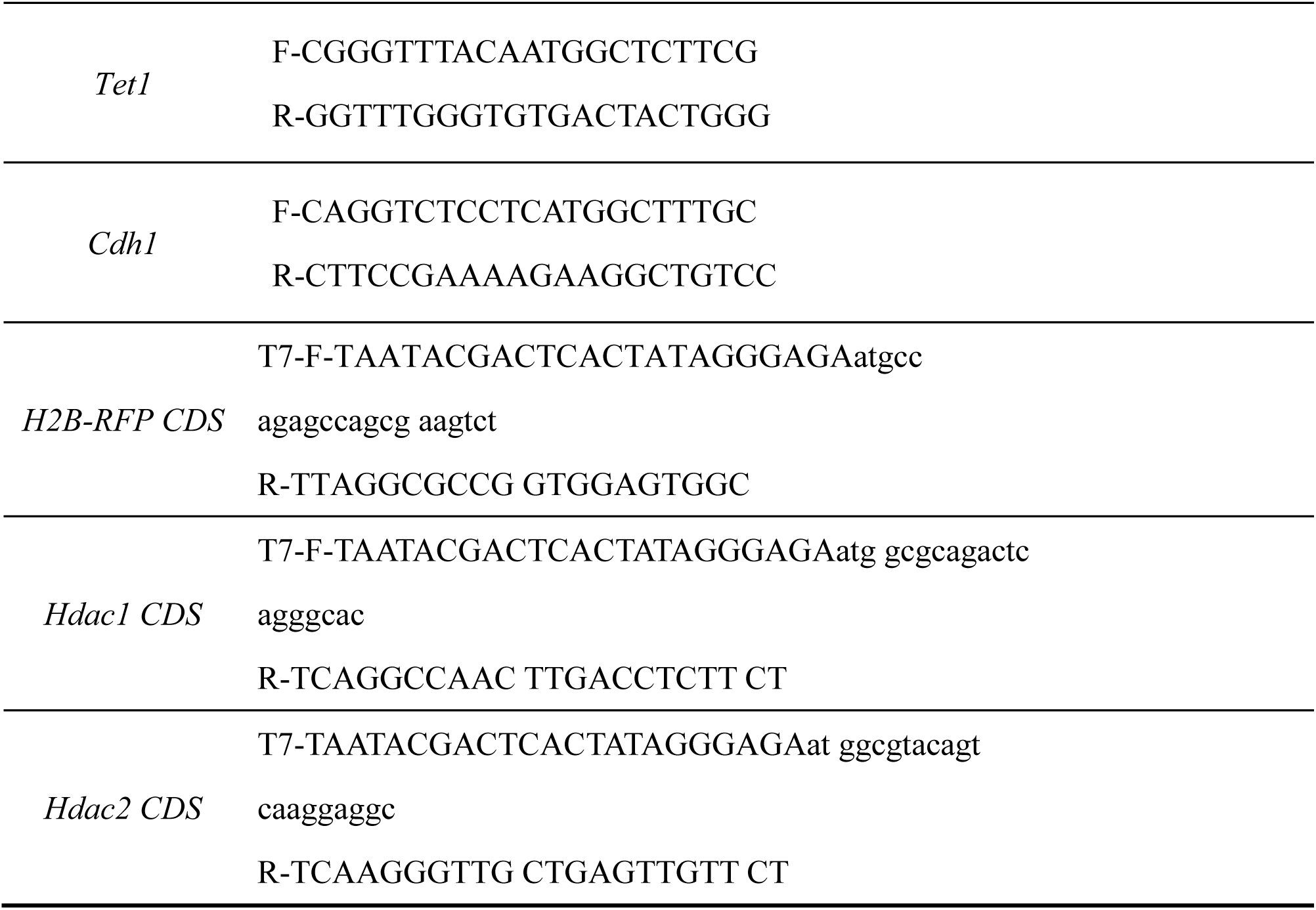
List of siRNAs and oligos used for qPCR and *in vitro* transcription

**Table S3.** List of antibodies used for immunofluorescence and Western Blot

| Target | Source | Dilution | Company | Catalogue |
| --- | --- | --- | --- | --- |
| Nanog | Rabbit | 1:200 | Cell Signaling Technology | 8822 |
| H3K4me3 | Rabbit | 1:200 | Cell Signaling Technology | 9751 |
| H3K9me2 | Rabbit | 1:200 | Cell Signaling Technology | 9753 |
| E-Cadherin | Rat | 1:1000 | Sigma | U3254 |
| DNMT3A | Mouse | 1:200 | Novus Biologicals | NB120-13888SS |
| DNMT3B | Mouse | 1:200 | Novus Biologicals | 52A1018 |
| TEAD4 | Mouse | 1:200 | Abcam | ab58310 |
| H4K5ac | Rabbit | 1:200 | Abcam | ab51997 |
| H3K14ac | Rabbit | 1:200 | Abcam | ab52946 |
| P-H3S10 | Rabbit | 1:1000 | Cell Signaling Technology | 3377 |
| 5mc | Mouse | 1:200 | Eurogentec | BI-MECY-1000 |
| 5-hmC | Rabbit | 1:200 | Active Motif | 39791 |
| HDAC2 | Rabbit | 1:200 | Abcam | ab32117 |
| α-Tubulin | Rabbit | 1:100 | Cell Signaling Technology | 2125 |
| HDAC1 | Mouse | 1:200 | Cell Signaling Technology | 5356 |
| CDX2 | Mouse | 1:200 | BioGenex | CDX2-88 |
| OCT3/4 | Mouse | 1:200 | Santa Cruz | sc-5279 |
| SOX2 | Rabbit | 1:50 | Millipore | AB5603 |
| YAP | Mouse | 1:200 | Santa Cruz | sc-101200 |
| GATA6 | Goat | 1:200 | R&D Systems | AF1700 |
| SOX17 | Goat | 1:200 | R&D Systems | AF1924 |
| Uhrf1 | Mouse | 1:200 | Santa Cruz | sc-373750 |
| P53-acetyl-lys379 | Rabbit | 1:200 | GeneTex | GTX88013 |
| β-actin | Mouse | 1:1000<br>(WB) | Beyotime | AF0003 |
| Donkey anti-Rabbit<br>594 |  | 1:1000 | Invitrogen | A21207 |
| Goat anti-Mouse<br>488 |  | 1:1000 | Invitrogen | A11001 |
| Donkey anti-Mouse<br>594 |  | 1:1000 | Invitrogen | A11032 |
| Donkey anti-Goat<br>594 |  | 1:1000 | Invitrogen | A11058 |
| Goat anti-Rat 488 |  | 1:1000 | Invitrogen | A11006 |
| Goat anti-Rabbit<br>488 |  | 1:1000 | Invitrogen | A11008 |
| Donkey anti-Mouse<br>488 |  | 1:1000 | Invitrogen | A21206 |

